# Decoding by Dynamics: Reframing Neural Decoding as Stable Control Inference with Behavior Priors

**DOI:** 10.64898/2026.07.25.740738

**Authors:** Zonghan Du, Lai Zhong Yuan, Liang Hu

## Abstract

Continuous neural decoding is fragile under nonstationary neural recordings because unconstrained sequence regressors can turn small mapping errors into temporally inconsistent and physically implausible motion. We propose **Neural State-Space Dynamic Movement Primitives (Neural SS-DMP)**, which shifts the inductive bias from the neural encoder to the decoded output space: instead of directly predicting kinematics, the model infers low-dimensional movement-primitive controls and realizes them through a differentiable second-order dynamical generator. This reframes decoding as structured control inference, shrinking the set of admissible trajectories while retaining expressivity through learned forcing inputs. Because a universal motor prior cannot capture subject-specific movement dynamics, we form a personalized generator by blending the base DMP dynamics with behavior-derived subject dynamics estimated solely from training kinematics. Across ECoG and multi-session spiking benchmarks, Neural SS-DMP improves strong offline baselines in accuracy, consistently improves trajectory smoothness, and shows slower degradation on chronologically held-out sessions under an offline window-causal protocol.

## 1 Introduction

Decoding continuous movement from neural activity is central to BCIs and systems neuroscience, yet remains fragile in realistic settings. Neural recordings are high-dimensional, noisy, and nonstationary [1], and limited evidence in a causal history window often makes the mapping ill-posed. Consequently, small perturbations—from noise or session-to-session drift—can cause not only pointwise errors but qualitatively distinct failures: when unconstrained, a decoder may emit temporally inconsistent, high-frequency motion that undermines usability.

Classic linear estimators are interpretable [2–4], and modern RNNs/Transformers are highly expressive [5–7], with recent alignment methods such as NoMAD[8] targeting cross-session robustness. However, we argue that fragility in ill-posed regimes is also an output-space problem: unconstrained sequence regression leaves too many trajectories feasible under limited evidence (Fig. 1). In such a large hypothesis space, mapping errors can appear as high-frequency artifacts; a decoder may be accurate on average yet dynamically inconsistent. This reframes robustness as a multi-level problem: beyond expressive neural-to-kinematic regression and neural-latent alignment under drift, we ask **how to constrain decoded behavior itself under limited and nonstationary neural evidence?**

**Figure 1:**
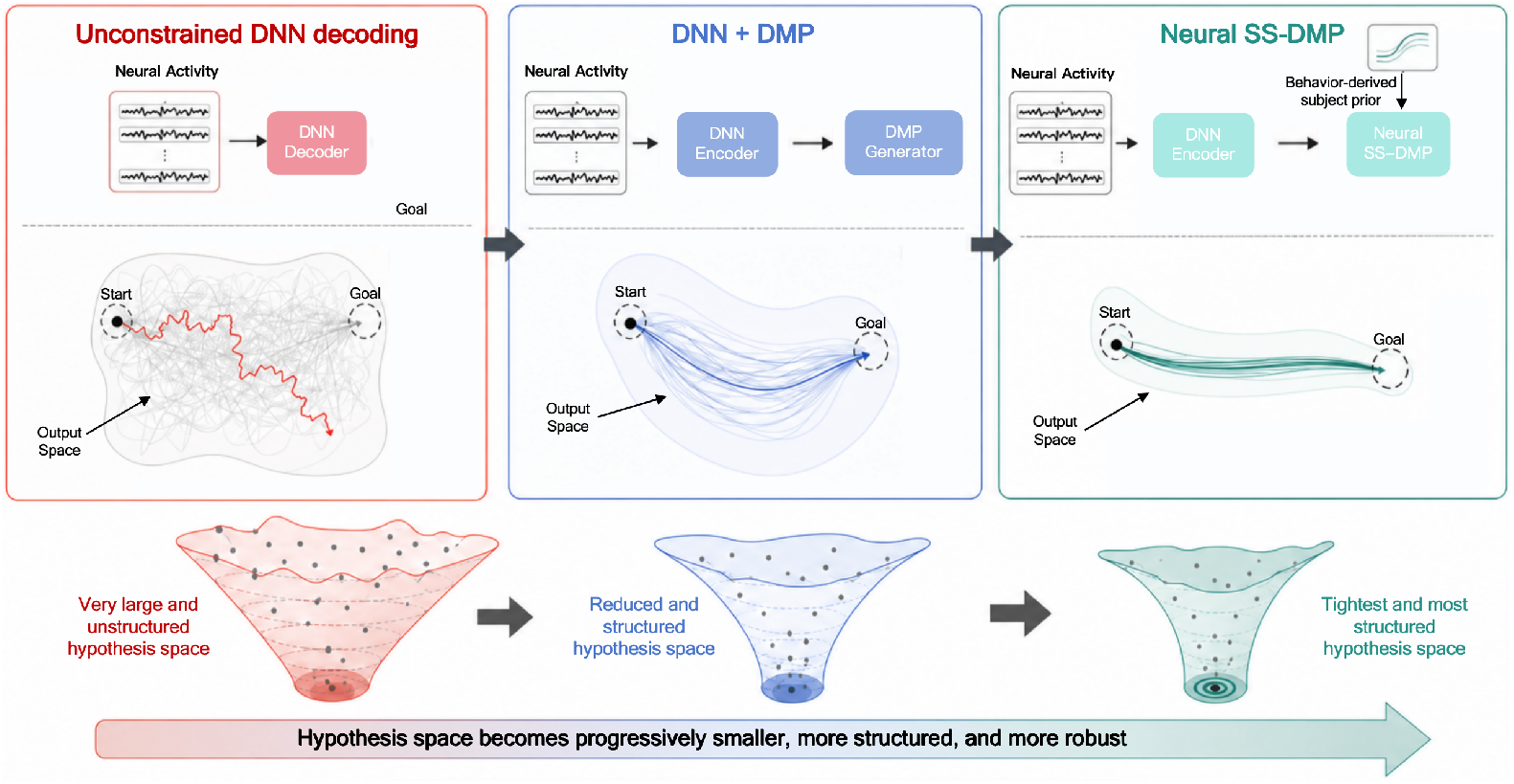
Motivation: progressively constraining the neural-to-kinematics hypothesis space. **Left:** Unconstrained DNN decoding leaves a large family of feasible trajectories. **Middle:** Adding a DMP (Dynamic Movement Primitives) generator restricts outputs to stable dynamical rollouts. **Right:** Neural SS-DMP further incorporates a behavior-derived subject prior, yielding a smaller, more structured, and more robust trajectory family.

This output-space perspective suggests drawing on motor control: decoded movement should be generated by structured dynamics, not emitted as an arbitrary sequence. Dynamic Movement Primitives (DMPs) formalize this idea by representing trajectories as stable attractors modulated by compact forcing functions [9]. Motivated also by the low-dimensional organization of neural population activity [10], we infer movement-primitive controls from neural evidence and realize them through an explicit dynamical generator. This complements neural-latent stabilization methods such as NoMAD, which seek stable neural coordinates under drift; our focus is to structure the behavioral trajectory family produced by the decoder.

We introduce **Neural SS-DMP**, a structured decoder that predicts primitive variables (goal, forcing, initialization) from a causal neural window and rolls them out through a differentiable state-space DMP. By restricting predictions to dynamically realizable trajectories, Neural SS-DMP suppresses high-frequency artifacts under mapping errors while retaining expressive time-varying shapes via learned forcing inputs. To avoid a one-size-fits-all motor prior, we blend the base DMP dynamics with behavior-derived subject dynamics estimated solely from training kinematics, providing personalization without held-out neural labels across heterogeneous feature types.

We evaluate Neural SS-DMP on two ECoG and two multi-session spiking datasets, comparing against classical, direct neural, and alignment-based baselines under the same within-window protocol. The model matches or improves offline decoding accuracy while producing smoother trajectories, and chronologically held-out analyses suggest slower degradation under session shift. Appendices J–K provide additional experimental results, analyses, and visualizations, among others.

### Contributions

- **Output-side robustness by construction**. Replace unconstrained regression with structured decoding: infer low-dimensional controls and roll them out via a stable state-space DMP.
- **Behavior-only personalization**. Blend subject-specific dynamics learned from training kinematics into the generator, enabling personalization without test-time kinematic labels.
- **Performance**. Show slower cross-session degradation on spiking benchmarks, while maintaining strong performance on ECoG.

## 2 Related Work

### Continuous neural decoding and state-space estimators

Continuous kinematic decoding in BCIs has classically used linear regression, Wiener filtering, and probabilistic state-space estimators such as Kalman filters, which are efficient and interpretable [11, 2–4, 12]. Another line of work models neural population activity with latent dynamical structure, including GPFA and LFADS, to capture low-dimensional neural dynamics [13–15]. Neural SS-DMP acts at a complementary level: rather than replacing latent neural-dynamics or alignment methods, it constrains the decoded behavior through a differentiable dynamical generator. Thus, neural-space methods that infer or stabilize latent population dynamics could in principle feed their latent states into SS-DMP primitive-control heads.

### Deep sequence decoders, ill-posed regression, and output-side structure

Modern decoders often use recurrent, convolutional, and attention-based sequence models to map neural time series to kinematics [1, 16, 7]. While expressive, direct sequence regression can be underconstrained when neural evidence is limited or nonstationary, allowing mapping errors to appear as temporally inconsistent, high-frequency artifacts [17, 18]. We address this failure mode by predicting compact primitive controls and generating trajectories through an explicit dynamical prior, restricting outputs to dynamically realizable rollouts.

### Movement primitives, differentiable dynamics, and personalization under drift

Movement primitives parameterize trajectories with stable dynamical systems, with DMPs providing a canonical formulation in motor control and robotics [19, 9]. Related machine-learning approaches use differentiable dynamical layers and simulators, including neural ODE-style formulations, for end-to-end learning with explicit state-space structure [20, 21]. Practical BCIs further require calibration or adaptation to inter-subject variability and nonstationarity [17, 22], and recent methods such as NoMAD target cross-session robustness by aligning neural representations with unlabeled neural data [8]. In contrast, Neural SS-DMP incorporates personalization as a behavior-only output-side prior: subject-specific stable dynamics are fit from training kinematics and blended into the generator, without supervised test-time recalibration.

## 3 Method

### 3.1 Overview

Neural SS-DMP is a **structured neural decoder** that performs control inference under an explicit dynamical prior. Given a window-causal neural history 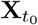 (ECoG raw/PSD or binned spikes), a Conv–Transformer produces embeddings (**z, z**_init_) (Fig. 2). MLP heads map them to primitive variables (**ĝ, ŵ**, **Ŷ**_0_, 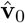), and a differentiable **SS-DMP** generator rolls out stable second-order dynamics (basis-function forcing) to output 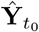. We inject a behavior-derived subject prior fit from **training kinematics only** by blending subject and base dynamics with fixed (*λ*_*A*_, *λ*_*B*_) = (0.2, 0.5), and train end-to-end by backpropagating a position loss (MSE) through the rollout. The generator targets smooth, goal-directed movements.

**Figure 2:**
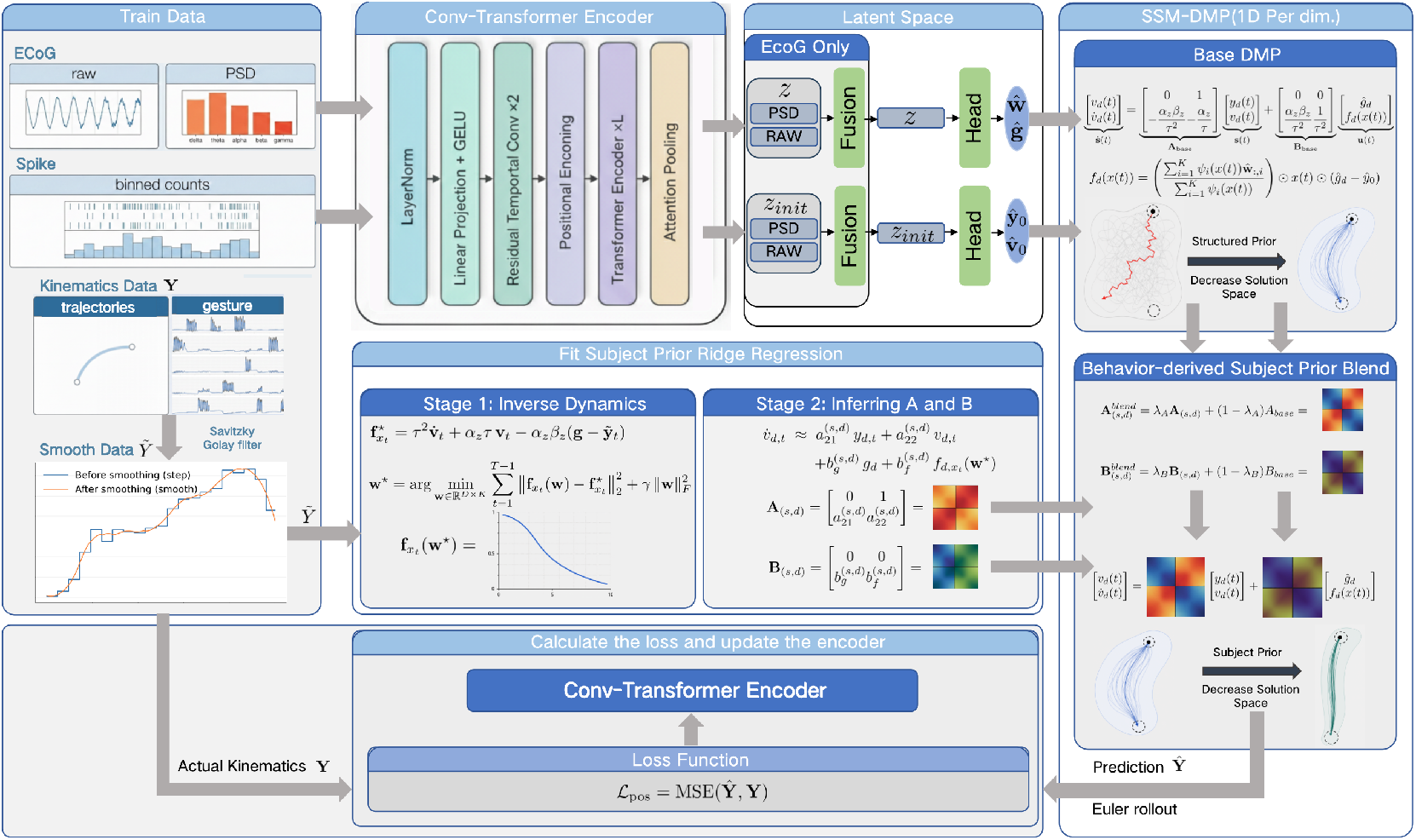
Neural SS-DMP: encoder-to-rollout with behavior-derived subject prior. A causal encoder maps the neural history window 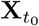 to latent summaries (**z, z**_init_); prediction heads output primitive parameters (**ĝ, ŵ**, **Ŷ**_0_, 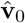), which are rolled out by the differentiable SS-DMP to generate kinematics. A subject-specific dynamical prior (*A*_(*s,d*)_, *B*_(*s,d*)_) is fit from training kinematics only (two-stage ridge) and blended with the base dynamics via fixed (*λ*_*A*_, *λ*_*B*_).

### 3.2 Problem: within-window decoding

Let **x**_*t*_ ∈ ℝ^*C*^ be neural features and **y**_*t*_ ∈ ℝ^*D*^ kinematics sampled every Δ*t*. Given a window of *H* neural samples starting at *t*_0_, 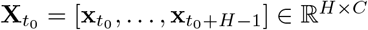, we predict the last *T* kinematic samples in the same window (*T* ≤ *H*). Define the target start as *t*_*s*_ = *t*_0_ + *H* − *T* and the target segment as 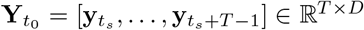. The offset (*H* − *T*)Δ*t* specifies where the target segment begins within the observed neural window. Neural SS-DMP starts the rollout(see Sec. 3.5) from an encoder-predicted initial state 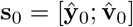 aligned to the target-segment start, and then rolls out for *T* steps to produce 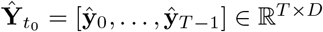, where 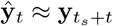 for *t* = 0, …, *T* − 1. See Appendix C for more details.

### 3.3 Neural encoder with within-aligned query pooling

A Conv–Transformer encoder maps the window-causal history (Sec. 3.2) to a control embedding **z** ∈ ℝ^*h*^ and an initialization embedding **z**_init_ ∈ ℝ^*h*^. Here **z** summarizes the neural evidence used to infer goal and forcing controls, while **z**_init_ is aligned to the start of the decoded target segment and is used to initialize the SS-DMP rollout. Full architectural details, including tokenization, convolution blocks, Transformer layers, dropout, and fusion MLPs, are provided in Appendix E.

#### Inputs and backbone

For ECoG, we use two modality streams over the same causal window: raw ECoG 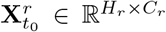 and PSD/band-power features 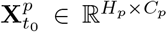, where *C*_*p*_ = *C*_*r*_*B* concatenates *B* frequency-band features per channel. These two views are both provided to the model and are encoded as parallel streams before fusion. For spiking data, we bin spike times with Δ*t*_bin_ to form 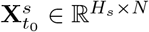 *N* units. Indexing modalities by *m* ∈ {*r, p, s*}, each stream 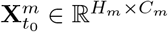 is processed independently by the same Conv–Transformer backbone to produce encoded tokens 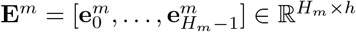. Self-attention is bidirectional within the observed window, but the protocol remains window-causal because no neural samples beyond *t*_0_ + *H* − 1 are accessed.

#### Within-aligned query pooling

The target segment starts at *t*_*s*_ = *t*_0_ + *H* − *T*, so the aligned raw-window index is *k*_*r*_ = *H* − *T* using 0-based indexing. For modality *m* with sequence length *H*_*m*_, we map this index to 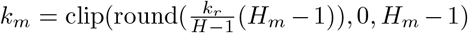. We take the aligned token as the initialization embedding, 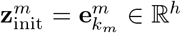, and obtain a global control embedding by using the same token as a length-1 query over the full encoded sequence:

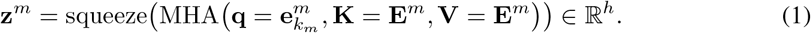

This separates local target-start information for initialization from full-window information for goal and forcing inference.

#### Multimodal fusion

For ECoG, raw and PSD streams are fused with lightweight MLPs: **z** = *ψ*([**z**^*r*^; **z**^*p*^]) and 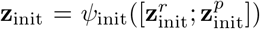, where [·; ·] denotes concatenation and *ψ, ψ*_init_ map the concatenated embeddings to ℝ^*h*^. For spiking data, we use the single spike stream directly: 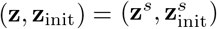

### 3.4 Primitive control heads

Let *D* be the kinematic dimension and *K* the number of DMP basis functions. Given (**z, z**_init_), lightweight readout heads predict the SS-DMP inputs: 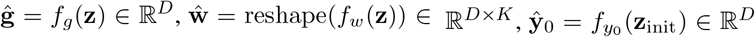 and 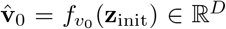. Here **ĝ** denotes the segment goal, i.e., the desired final position at the end of the within-window target segment. The weight matrix **ŵ** contains basis weights, where **ŵ** _*d,i*_ is the weight of basis function *i* for kinematic dimension *d*. The generator is initialized as 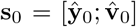, where **Ŷ**_0_ and 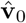 denote the initial position and velocity state, respectively.

### 3.5 SS-DMP generator: from DMP to state-space

Throughout this section, scalar equations are written for one kinematic dimension, while bold symbols denote stacked *D*-dimensional quantities. Continuous-time variables are written with parentheses, e.g., *y*(*t*) or **s**_*d*_(*t*), whereas sampled rollout values use subscripts, e.g., *y*_*d,t*_ or **s**_*d,t*_.

#### Canonical DMP transformation system (per dimension)

Given predicted primitive variables (**ŵ**, **ĝ**) and an initial state (**Ŷ**_0_, 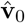), the **State-Space DMP (SS-DMP)** deterministically produces a length-*T* kinematic segment by rolling out a second-order attractor-like dynamics with step size Δ*t* (Full generator details are provided in Appendix F.). For a single output dimension, the continuous-time DMP transformation system is

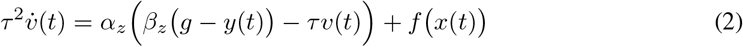

where 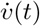 is acceleration, *v*(*t*) is velocity, *y*(*t*) is position, *g* ∈ ℝ is the goal, *τ* is time constant, *α*_*z*_, *β*_*z*_ *>* 0 are stiffness/damping gains. We apply this form independently across the *D* output dimensions and write the stacked state-space form below.

#### Phase variable

Let *x*_*t*_ ∈ ℝ denote the scalar phase, distinct from neural features **x**_*t*_. With *x*(0) = 1, the canonical system 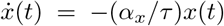 admits the closed-form discrete update *x*_*t*+1_ = exp(−*α*_*x*_Δ*t/τ*)*x*_*t*_, where *α*_*x*_ *>* 0 is the phase decay rate.

#### Forcing function parameterization

We use *K* Gaussian radial basis functions (RBFs) *ψ*_*i*_(*x*(*t*)) = exp (− *h*_*i*_(*x*(*t*) − *c*_*i*_)^2^, with centers placed to uniformly cover the *T* -step segment (in time) and mapped into phase *x*, and widths set from adjacent center spacing to ensure smooth overlap (Appendix F.1). Given **ŵ** ∈ ℝ^*D×K*^, the continuous-time forcing(**f** (*t*) ∈ ℝ^*D*^) input is

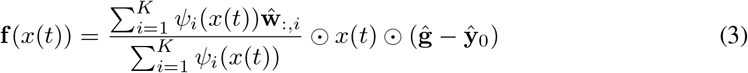

#### State-space form and rollout

For each kinematic dimension *d* = 1, …, *D*, define the state **s**_*d*_(*t*) = [*y*_*d*_(*t*); *v*_*d*_(*t*)] ∈ ℝ^2^ with 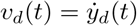. The continuous-time SS-DMP dynamics are

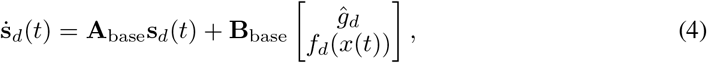

where 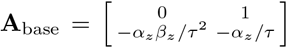 and 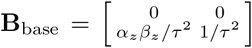. Here *α*_*z*_ and *β*_*z*_ are the DMP stiffness and damping gains, and *τ* is the temporal scaling parameter. We discretize Eq.(4) with a forward Euler step of size Δ*t*:

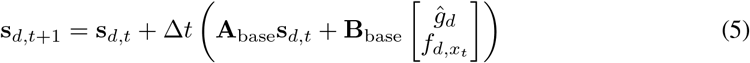

We start the rollout from the encoder-predicted initial state 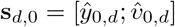 and record positions for *t* = 0, …, *T* − 1, producing 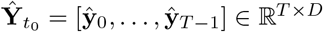 (Sec. 3.2).

### 3.6 Behavior-derived subject priors for SS-DMP dynamics

To capture individual motor characteristics and further restrict the solution space, we learn a subject-specific dynamics prior from training kinematics only (Details of the prior-fitting procedure are provided in Appendix G.). We fit independent second-order dynamics per kinematic dimension. For subject *s* and dimension *d* ∈ {1, …, *D*}, with state **s**_*d*_(*t*) = [*y*_*d*_(*t*); *v*_*d*_(*t*)] ∈ ℝ^2^ and inputs (*g*_*d*_, *f*_*d*_(*t*)), where **A**_(*s,d*)_, **B**_(*s,d*)_ ∈ ℝ^2*×*2^:

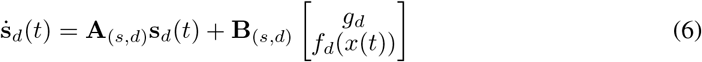

#### Stage 1: infer forcing from training behavior

We infer discrete forcing targets 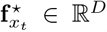 by evaluating the rearranged continuous-time DMP equation (Eq. 2) dimension-wise at sampled training steps, using finite-difference velocity and acceleration estimates from training kinematics. To stabilize these derivative estimates, we use the next sample **y**_*T*_ only for finite differences and smooth kinematics only when forming derivative targets; details are provided in Appendix D.5. Because the SS-DMP generator realizes forcings only in the RBF family of Eq. (3), we project 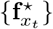 onto this family by fitting **w**^⋆^ ∈ ℝ^*D×K*^ via ridge regression:

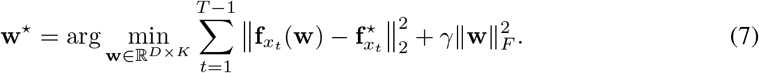

We use a fixed ridge penalty *γ* = 10^−3^ and use the fitted forcing values **f**_*x*_ (**w**^⋆^) for Stage 2.

#### Stage 2 (stable prior): regress (A_(*s,d*)_, B_(*s,d*)_)

Using the training states and generator-consistent forcing values constructed in Stage 1, we form tuples (*y*_*d,t*_, *v*_*d,t*_, *g*_*d*_, *f*_*d,x*_ (**w**^⋆^)) and fit continuous-time coefficients independently per dimension:

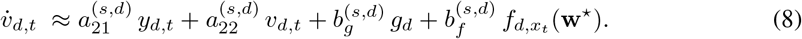

We then assemble the subject-specific dynamics as 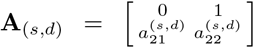 and 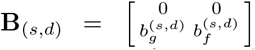. To ensure stability, we enforce tr(**A**_(*s,d*)_) *<* 0 and det(**A**_(*s,d*)_) *>* 0 (stable second-order system) via constrained ridge regression in all main experiments.

#### Injecting the prior (fixed blending)

During decoding, we blend the base SS-DMP dynamics matrices **A**_base_ and **B**_base_ defined in Sec. 3.5 with the learned subject-specific prior using separate fixed coefficients:

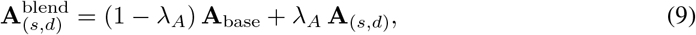

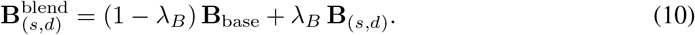

Here *λ*_*A*_ personalizes intrinsic dynamics (effective stiffness/damping) and *λ*_*B*_ personalizes input gains. We use fixed defaults (e.g., *λ*_*A*_=0.2, *λ*_*B*_=0.5 unless stated) and do not use validation/test kinematics for fitting or calibration.

## 4 Experiments

### 4.1 Datasets

We evaluate Neural SS-DMP on two human ECoG datasets and two intracortical spiking datasets (Tab. 1). All datasets provide continuous kinematics and use the within-window protocol (Sec. 3.2); dataset-specific preprocessing and Δ*t* are in Appendix Tab. 6. See Appendix D.1 for more details.

### 4.2 Protocol, splits, and inputs

For ECoG, we use raw+PSD fusion (Sec. 3.3). Raw ECoG is common-average referenced and offline filtered with a 1–200 Hz bandpass and 60 Hz notch filtering to form 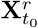. PSD features 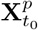 are computed from the same history window using Welch band-power features and summarized in 25 non-overlapping 4 Hz bands spanning 1–100 Hz. Raw time-domain ECoG features are z-scored using training-set statistics, while PSD features use the window-local, label-free normalization. For spiking data, we bin spike counts at Δ*t*_bin_ = Δ*t*, apply causal exponential smoothing, and z-score using held-in training-set statistics. All preprocessing and normalization choices are applied identically to all models using the same input modality. See Appendix D.2, D.3 and D.4 for more details.

### 4.3 Baselines, training, and metrics

We compare against a Kalman filter and direct sequence decoders (Transformer, TCN [27], RNN, LSTM) under the same protocol, splits, targets, and preprocessing. On multi-session spiking datasets only, we additionally evaluate NoMAD [8], trained on labeled held-in sessions and performing unlabeled test-session alignment using neural activity only (no test-session behavioral labels). Details regarding the reproduction of all baseline models are provided in Appendix I. All neural models are trained with the same segment-wise position MSE on their trajectory outputs, ℒ = MSE(**Ŷ**, **Y**) (Appendix H). Unless stated otherwise, we use *α*_*z*_=25, *β*_*z*_=6.25, *α*_*x*_=1, *τ* ≡ 1 (physical time via Δ*t*), *K*=10 RBF bases, and fixed prior blending (default *λ*_*A*_ = 0.2, *λ*_*B*_ = 0.5), selected once using held-out validation data and then kept fixed for all test evaluations (Appendix J.9). We train with AdamW (lr 3 ×10^−4^, wd 10^−2^), cosine decay with 5% warmup, batch size 64, and a common budget (max 200 epochs with early stopping on validation loss, patience 20); we select the checkpoint with lowest validation loss. We report mean ± 95% CIs over 5 random seeds (Appendices B.1 and B.2); model-specific widths, including embedding/hidden sizes and layers/heads, are provided in Appendix E. We report variance-weighted *R*^2^, mean Pearson correlation, RMSE in original units after undoing any target standardization, and mean squared jerk (MSJ; Appendix B.2).

### 4.4 Results

#### 4.4.1 Main comparison across datasets

Table 2 summarizes within-window decoding under the unified protocol of Sec. 3.2 with (*H*_sec_, *T*_sec_) = (0.7 s, 0.5 s). We report variance-weighted *R*^2^, mean Pearson correlation (Corr), RMSE, and mean squared jerk (MSJ; lower is smoother). All neural models use identical splits, preprocessing, and evaluation windows, and are selected by validation loss; the Kalman filter is fit on the training split. For MouseTrack and BCI IV-4, metrics are averaged across subjects within each seed before reporting mean ± 95% CI across 5 random seeds. NoMAD is evaluated only on the spiking datasets, where its standard unlabeled test-session alignment setting is applicable. Across the four benchmarks, Neural SS-DMP achieves the best variance-weighted *R*^2^ and Corr on every dataset. It also obtains the lowest RMSE on three of four datasets, with NoMAD slightly lower on H1. Most consistently, Neural SS-DMP yields the lowest MSJ in every main comparison, often by a large margin relative to direct sequence decoders. These results indicate that the proposed output-side dynamical generator improves the accuracy–smoothness trade-off: it produces substantially smoother trajectories while preserving or improving offline decoding accuracy across ECoG and spiking benchmarks. Additional analyses, results and visualizations are provided in Appendices J and K.

**Table 1:**
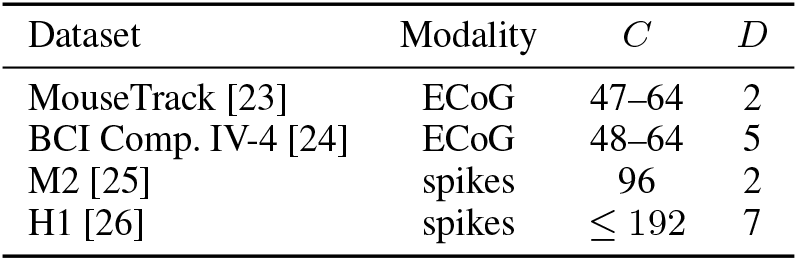
Dataset summary. *C* is the number of ECoG channels or spike units; *D* is the kinematic dimensionality.

**Table 2:** Main results (within-window). Variance-weighted *R*^2^ and Corr (higher is better), RMSE and MSJ (lower is better) for *H*_sec_=0.7 s and *T*_sec_=0.5 s. MouseTrack and BCI IV-4 averages over subjects within each seed. Entries are mean ± 95% CI across 5 random seeds. For ECoG datasets, all models use matched raw+PSD features; NoMAD is evaluated on spiking datasets only (M2, H1) under the same within window protocol; ECoG entries are not applicable (“–”).

| Dataset | Metric | Ours | Baselines |  |  |  |  |  |
| --- | --- | --- | --- | --- | --- | --- | --- | --- |
|  |  | Neural SS-DMP | Transformer | TCN | LSTM | RNN | Kalman | NoMAD |
| M2<br>(spikes) | $R^2 \uparrow$ | **0.44 ± 0.08** | 0.31 ± 0.10 | 0.36 ± 0.06 | 0.28 ± 0.09 | 0.38 ± 0.09 | 0.26 ± 0.07 | 0.41 ± 0.09 |
| | Corr ↑ | **0.63 ± 0.11** | 0.50 ± 0.12 | 0.52 ± 0.07 | 0.55 ± 0.06 | 0.48 ± 0.13 | 0.41 ± 0.09 | 0.58 ± 0.10 |
| | RMSE ↓ | **0.70 ± 0.16** | 0.93 ± 0.22 | 0.82 ± 0.14 | 1.10 ± 0.28 | 0.97 ± 0.17 | 1.04 ± 0.19 | 0.78 ± 0.18 |
| | MSJ ↓ | **1.62 ± 0.10** | 5.47 ± 0.46 | 3.95 ± 0.31 | 4.89 ± 0.41 | 2.87 ± 0.23 | 3.45 ± 0.27 | 2.95 ± 0.25 |
| H1<br>(spikes) | $R^2 \uparrow$ | **0.26 ± 0.03** | 0.20 ± 0.05 | 0.25 ± 0.02 | 0.18 ± 0.06 | 0.23 ± 0.03 | 0.12 ± 0.07 | **0.26 ± 0.04** |
| | Corr ↑ | **0.42 ± 0.06** | 0.39 ± 0.04 | 0.31 ± 0.08 | 0.36 ± 0.05 | 0.34 ± 0.07 | 0.26 ± 0.10 | 0.36 ± 0.06 |
| | RMSE ↓ | 1.12 ± 0.15 | 1.29 ± 0.27 | 1.20 ± 0.18 | 1.14 ± 0.11 | 1.26 ± 0.21 | 1.47 ± 0.33 | **1.08 ± 0.16** |
| | MSJ ↓ | **1.14 ± 0.09** | 3.88 ± 0.33 | 2.79 ± 0.21 | 3.21 ± 0.27 | 2.12 ± 0.15 | 2.94 ± 0.18 | 2.35 ± 0.18 |
| BCI IV-4<br>(ECoG) | $R^2 \uparrow$ | **0.34 ± 0.09** | 0.33 ± 0.05 | 0.27 ± 0.11 | 0.30 ± 0.08 | 0.24 ± 0.10 | 0.21 ± 0.07 | – |
| | Corr ↑ | **0.54 ± 0.11** | 0.46 ± 0.14 | 0.51 ± 0.09 | 0.47 ± 0.12 | 0.53 ± 0.07 | 0.40 ± 0.10 | – |
| | RMSE ↓ | **0.82 ± 0.07** | 0.91 ± 0.12 | 0.84 ± 0.06 | 1.02 ± 0.15 | 0.95 ± 0.09 | 1.11 ± 0.18 | – |
| | MSJ ↓ | **0.97 ± 0.07** | 4.03 ± 0.38 | 3.26 ± 0.29 | 3.84 ± 0.33 | 2.35 ± 0.19 | 3.17 ± 0.21 | – |
| MouseTrack<br>(ECoG; avg.) | $R^2 \uparrow$ | **0.42 ± 0.06** | 0.35 ± 0.10 | 0.39 ± 0.05 | 0.41 ± 0.04 | 0.33 ± 0.12 | 0.28 ± 0.07 | – |
| | Corr ↑ | **0.63 ± 0.12** | 0.58 ± 0.10 | 0.62 ± 0.08 | 0.55 ± 0.13 | 0.60 ± 0.09 | 0.47 ± 0.11 | – |
| | RMSE ↓ | **0.72 ± 0.08** | 0.73 ± 0.07 | 0.86 ± 0.14 | 0.79 ± 0.09 | 0.88 ± 0.16 | 1.03 ± 0.20 | – |
| | MSJ ↓ | **2.08 ± 0.14** | 6.12 ± 0.57 | 4.48 ± 0.34 | 5.23 ± 0.39 | 3.45 ± 0.28 | 4.01 ± 0.33 | – |

#### 4.4.2 Qualitative M2 reconstructions

Figure 3a visualizes M2 reconstructions from Neural SS-DMP, NoMAD, and RNN under identical within-window evaluation splits. To avoid cherry-picking, we rank non-overlapping evaluation windows by Neural SS-DMP variance-weighted *R*^2^ and show a contiguous interval containing the median-ranked window by tiling adjacent non-overlapping predictions. Neural SS-DMP follows the target level and transition timing while producing temporally coherent rollouts; NoMAD captures the coarse trajectory but shows larger local deviations, and RNN exhibits larger drift and high-frequency fluctuations in this segment. Additional qualitative reconstructions are provided in Appendix J.1.

**Figure 3:**
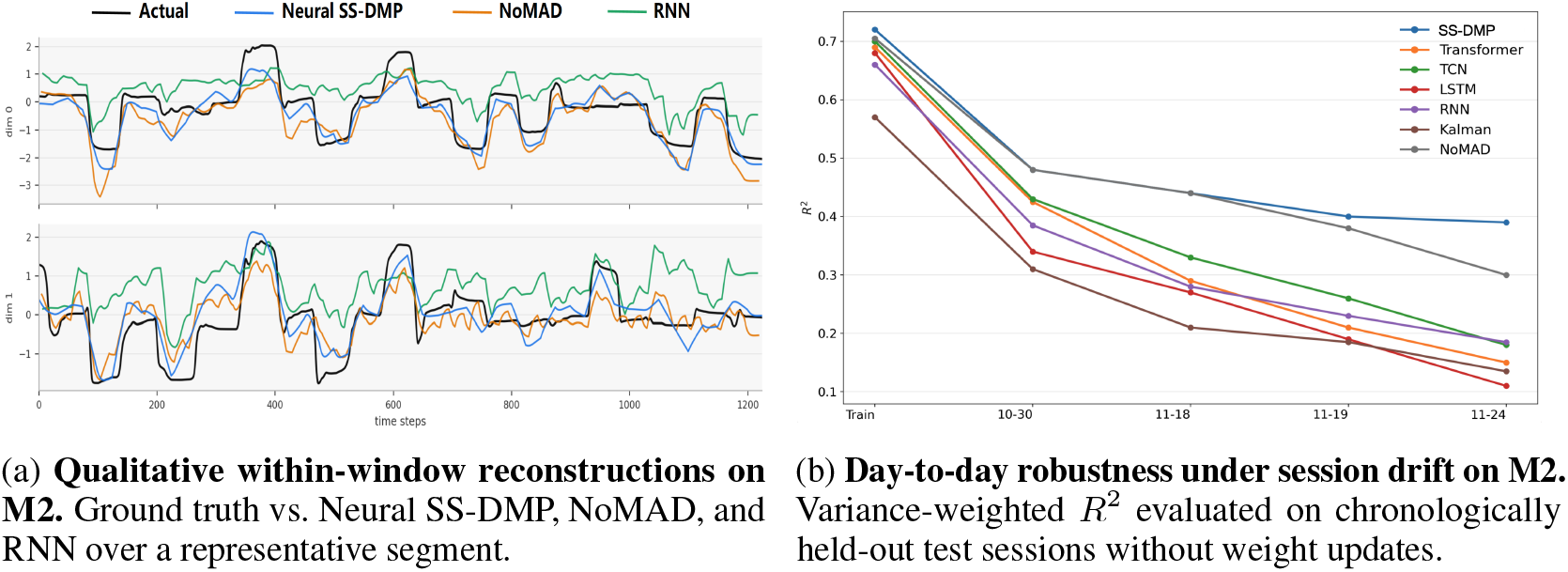
Qualitative reconstruction and session-drift robustness on M2 (spikes).

#### 4.4.3 Robustness to day-to-day shift (session drift)

We probe day-to-day distribution shift on the multi-session spiking benchmark using **M2**, whose chronologically held-out (HO) test files provide recording dates, enabling performance tracking versus increasing time from training. Under the within window protocol, we train a single model on the HI split and evaluate each dated HO session independently in chronological order, so across-session changes primarily reflect drift rather than supervised adaptation. Figure 3b reports variance-weighted *R*^2^: the leftmost point summarizes HI-session performance under the same procedure and subsequent points correspond to dated HO sessions; curves are means over 5 random seeds. Neural SS-DMP maintains higher *R*^2^ and degrades more slowly than sequence-regression baselines and NoMAD, indicating improved robustness to session drift under decoding.

#### 4.4.4 Smoothing, projection, and prior ablations

Table 3 isolates the contributions of output-side structure and behavior-derived dynamics. We compare the full Neural SS-DMP against four diagnostic settings under the same within-window protocol: a direct Transformer regressor without structured rollout, Transformer+EMA with causal post-hoc smoothing, Regress+Project with post-hoc projection into the SS-DMP trajectory family, and SS-DMP w/o prior, which removes subject-specific dynamics blending. Post-hoc smoothing substantially reduces trajectory roughness, but does not recover the accuracy of end-to-end Neural SS-DMP. For example, on M2, Transformer+EMA reduces MSJ from 5.47 to 1.54, slightly below the full model’s 1.62, but its *R*^2^ remains much lower (0.33 vs. 0.44). Regress+Project also improves smoothness relative to direct regression and partially improves accuracy, but remains below end-to-end SS-DMP in *R*^2^ across all datasets. Removing the behavior-derived prior consistently lowers *R*^2^ and increases MSJ relative to the full model on all four benchmarks. Together, these ablations show that the final accuracy–smoothness trade-off comes from both end-to-end structured control inference and personalized dynamics, rather than from ordinary smoothing or post-hoc projection alone.

**Table 3:** Ablations on smoothing, post-hoc projection, and dynamics personalization. We report variance-weighted *R*^2^ (higher is better) and mean squared jerk (MSJ; lower is better) under the same within-window protocol as Table 2. Transformer+EMA (Exponential Moving Average) applies causal exponential moving-average smoothing to Transformer predictions at test time. Regress+Project fits SS-DMP primitives to a direct regression output and re-rolls the generator. SS-DMP w/o prior removes behavior-derived subject-specific dynamics blending.

| Method | $R^2 \uparrow$ | | | | MSJ ↓ | | | |
| --- | --- | --- | --- | --- | --- | --- | --- | --- |
|  | M2 | H1 | BCI IV-4 | MouseTrack | M2 | H1 | BCI IV-4 | MouseTrack |
| Transformer | 0.31 ± 0.10 | 0.20 ± 0.05 | 0.33 ± 0.05 | 0.35 ± 0.10 | 5.47 ± 0.46 | 3.88 ± 0.33 | 4.03 ± 0.38 | 6.12 ± 0.57 |
| Transformer+EMA | 0.33 ± 0.09 | 0.21 ± 0.05 | 0.32 ± 0.06 | 0.36 ± 0.09 | 1.54 ± 0.12 | 1.32 ± 0.11 | 1.09 ± 0.08 | 2.31 ± 0.18 |
| Regress+Project | 0.34 ± 0.09 | 0.22 ± 0.05 | 0.32 ± 0.07 | 0.38 ± 0.09 | 1.88 ± 0.13 | 1.42 ± 0.10 | 1.26 ± 0.09 | 2.55 ± 0.17 |
| SS-DMP w/o prior | 0.37 ± 0.09 | 0.24 ± 0.04 | 0.33 ± 0.09 | 0.40 ± 0.07 | 1.74 ± 0.12 | 1.23 ± 0.08 | 1.08 ± 0.08 | 2.20 ± 0.15 |
| SS-DMP full | **0.44 ± 0.08** | **0.26 ± 0.03** | **0.34 ± 0.09** | **0.42 ± 0.06** | **1.62 ± 0.10** | **1.14 ± 0.09** | **0.97 ± 0.07** | **2.08 ± 0.14** |

#### 4.4.5 SS-DMP as a plug-in output layer

Table 4 tests whether the benefit of Neural SS-DMP comes from the output-side dynamical generator rather than from a particular encoder backbone. For each backbone, we compare direct trajectory regression with a plug-in SS-DMP output layer: the direct model predicts kinematics **Ŷ** pointwise, whereas the SS-DMP variant predicts primitive variables (**ĝ, ŵ**, **Ŷ**_0_, 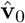) and generates trajectories through the differentiable rollout. All variants use the same datasets, splits, preprocessing, target windows, and training budget as Table 2. Across all four backbones and all four datasets, replacing the direct output head with SS-DMP improves variance-weighted *R*^2^ and substantially reduces MSJ. This consistent pattern supports the interpretation that the gain is an output-side effect of structured dynamical generation, rather than a consequence of the Conv–Transformer encoder alone.

**Table 4:** SS-DMP as a plug-in output layer across encoder backbones. For each backbone, we compare direct trajectory regression with replacing the direct output head by SS-DMP primitive-control heads. We report variance-weighted *R*^2^ (higher is better) and mean squared jerk (MSJ; lower is better) under the same within-window protocol as Table 2.

| Backbone | Output layer | $R^2 \uparrow$ | | | | MSJ ↓ | | | |
| --- | --- | --- | --- | --- | --- | --- | --- | --- | --- |
|  |  | M2 | H1 | BCI IV-4 | MouseTrack | M2 | H1 | BCI IV-4 | MouseTrack |
| Conv-Transformer | Direct | 0.31 ± 0.10 | 0.20 ± 0.05 | 0.33 ± 0.05 | 0.35 ± 0.10 | 5.47 ± 0.46 | 3.88 ± 0.33 | 4.03 ± 0.38 | 6.12 ± 0.57 |
|  | +SS-DMP | **0.44 ± 0.08** | **0.26 ± 0.03** | **0.34 ± 0.09** | **0.42 ± 0.06** | **1.62 ± 0.10** | **1.14 ± 0.09** | **0.97 ± 0.07** | **2.08 ± 0.14** |
| TCN | Direct | 0.36 ± 0.06 | 0.25 ± 0.02 | 0.27 ± 0.11 | 0.39 ± 0.05 | 3.95 ± 0.31 | 2.79 ± 0.21 | 3.26 ± 0.29 | 4.48 ± 0.34 |
| | +SS-DMP | 0.42 ± 0.07 | **0.26 ± 0.03** | 0.33 ± 0.09 | **0.42 ± 0.06** | 1.70 ± 0.12 | 1.18 ± 0.08 | 1.02 ± 0.07 | 2.14 ± 0.13 |
| LSTM | Direct | 0.28 ± 0.09 | 0.18 ± 0.06 | 0.30 ± 0.08 | 0.41 ± 0.04 | 4.89 ± 0.41 | 3.21 ± 0.27 | 3.84 ± 0.33 | 5.23 ± 0.39 |
| | +SS-DMP | 0.39 ± 0.08 | 0.24 ± 0.04 | 0.33 ± 0.08 | 0.42 ± 0.05 | 1.80 ± 0.14 | 1.26 ± 0.09 | 1.07 ± 0.08 | 2.22 ± 0.16 |
| RNN | Direct | 0.38 ± 0.09 | 0.23 ± 0.03 | 0.24 ± 0.10 | 0.33 ± 0.12 | 2.87 ± 0.23 | 2.12 ± 0.15 | 2.35 ± 0.19 | 3.45 ± 0.28 |
| | +SS-DMP | 0.41 ± 0.08 | 0.25 ± 0.03 | 0.32 ± 0.09 | 0.40 ± 0.07 | 1.75 ± 0.13 | 1.22 ± 0.08 | 1.05 ± 0.07 | 2.18 ± 0.15 |

## 5 Conclusion, Limitations and Broader Impact

Neural SS-DMP reframes continuous decoding as control inference under an explicit dynamical prior, inferring compact primitive variables realized through stable differentiable rollout. This output-space constraint suppresses high-frequency artifacts and temporal inconsistencies without relying on post-hoc smoothing. Control-space analyses show compact, anisotropic inferred controls, supporting structured control learning. To handle subject variability, we blend a behavior-derived stable dynamics prior into the generator, providing empirical-Bayes-style shrinkage from training kinematics only. Across ECoG and multi-session spiking benchmarks, Neural SS-DMP improves offline decoding accuracy and smoothness while suggesting slower degradation on chronologically held-out M2 sessions. However our evaluation is offline and window-causal rather than closed-loop or co-adaptive, so future work should test whether the same output-side constraints improve online BCI control under user feedback. The current SS-DMP prior is also tailored to continuous movement trajectories; discontinuous, or non-movement outputs may require different task-specific generators. More broadly, more stable neural decoding could support smoother assistive BCI control, but deployment requires safeguards for neural-data privacy, overinterpretation of decoded intent, and premature clinical use before closed-loop validation.

## A Technical Appendix

The appendix is organized to make reproducibility, implementation details, experimental controls, and supplementary evidence easy to locate. Table 5 provides clickable links from common reviewer questions to the exact appendix sections and related main-text claims.

**Table 5:** Appendix roadmap. Reviewer-facing guide to reproducibility, implementation details, baseline controls, supplementary analyses, and interpretability results.

| Reviewer-facing guide | Appendix location | Main-text location |
| --- | --- | --- |
| **Reproducibility and evaluation protocol** |  |  |
| How are seeds, metrics, CIs, error bars, and compute defined? | App. B: Reproducibility (seeds, metrics/CIs, compute) | Secs. 4.3 |
| How are windows, targets, stride, splits, and valid masks constructed? | App. C: Windowing, Splits, and Input Construction | Sec. 3.2 |
| What are the dataset-specific preprocessing and sampling details? | App. D: Datasets and Preprocessing (ECoG raw/PSD, spikes, targets/velocity) | Secs. 4.1–4.2; Tab. 1 |
| How is chronological held-out/session-drift evaluation performed? | App. C.2: Spike splits and masks; App. D.4: Spike preprocessing | Sec. 4.4.3; Fig. 3b |
| **Model implementation** |  |  |
| How is the neural encoder implemented? | App. E: Encoder Architecture (raw+PSD streams, spike stream, tokenization, Conv–Transformer, query pooling, fusion) | Sec. 3.3; Fig. 2 |
| How is the SS-DMP generator implemented? | App. F: SS-DMP Generator (phase, RBF basis, forcing, Euler rollout, prior injection) | Sec. 3.5; Eqs. 3–5 |
| How is the behavior-derived subject prior fit and injected? | App. G: Behavior-Derived Priors (Stage 1 forcing projection, Stage 2 stable ridge, fixed blending, sanity checks) | Sec. 3.6; Eqs. 7–10 |
| How are the loss, optimizer, early stopping, and model selection defined? | App. H: Loss and Optimization | Sec. 4.3 |
| **Baselines and fairness controls** |  |  |
| How are Kalman, neural baselines, and NoMAD implemented fairly? | App. I: Baselines and Fairness (Kalman, neural baselines, NoMAD) | Sec. 4.3; Tab. 2 |
| How are post-hoc projection and smoothing controls implemented? | App. I.4: Transformer+EMA; App. I.5: Regress+Project | Sec. 4.4.4; Tab. 3 |
| **Supplementary results, ablations, and robustness checks** |  |  |
| Where are qualitative reconstructions? | App. J.1: Extended Results | Sec. 4.4.2; Fig. 3a |
| Where are the planned statistical tests for the main accuracy comparisons? | App. J.2: Paired Permutation Tests | Sec. 4.4.1; Tab. 2 |
| Why use a DMP generator rather than an unconstrained sequence output or ordinary smoothing? | App. J.3: Controlled Mechanistic Validation of DMP Trajectory Generation | – |
| Does the structured-dynamics advantage persist without independent window resets? | App. J.4: Sequential No-Reset Evaluation | – |
| Where are additional metrics and history-length sensitivity analyses? | App. J.5: Additional Metrics; App. J.6: History-Length Sweep | – |
| Where are analyses of the learned subject dynamics? | App. J.7: Learned Subject-Dynamics Priors | – |
| Does raw+PSD fusion help over raw-only or PSD-only ECoG inputs? | App. J.8: ECoG Feature Ablation | – |
| How sensitive is the method to prior strength? | App. J.9: Prior-Strength Sweep | – |
| Does the behavior-derived subject prior remain useful under velocity-distribution shift? | App. J.10: Velocity-Shift Stress Test | – |
| **Interpretability and control-space analysis** |  |  |
| What structure appears in the inferred forcing/control space? | App. K: Forcing Geometry (compactness, PCA planes/rays, 2D projections, takeaway) | – |

## B Reproducibility (Implementation-Exact)

### B.1 Seeds and Determinism

All reported numbers are averaged over *five* independent runs that differ only in the random seed (affecting model initialization, dropout, and training-sample order). Concretely:

- **Training/evaluation seeds**. We run five independent trainings with distinct random seeds, and evaluate each trained model on the same fixed data splits.
- **Visualization subsampling seed**. For forcing-geometry visualizations (Appendix K; Fig. 11), we fix a separate random seed so that the subsampled evaluation windows used for plotting are identical across runs.
- **Determinism note**. We do not claim bitwise determinism across all hardware/software stacks: GPU kernels, parallelism, and data loading can introduce minor nondeterminism. Reproducibility is therefore quantified via uncertainty across seeds (Appendix. B.2).
- **Leakage prevention**. All train/validation/test splits are constructed prior to training and remain fixed across seeds; for contiguous temporal holdouts, the three splits are disjoint in time.

### B.2 Metrics and 95% Confidence Intervals

This section fixes *exactly* the metric and uncertainty definitions used in Sec. 4.4.1 (Tab. 2) and in the robustness plot (Fig. 3b).

#### Notation

Let **Y** ∈ ℝ^*N×T ×D*^ and **Ŷ** ∈ ℝ^*N×T ×D*^ denote the ground-truth and predicted kinematic segments over *N* evaluation windows, each of horizon length *T* and output dimension *D*. We flatten over windows and time to obtain **Y**_flat_, **Ŷ** _flat_ ∈ ℝ^(*NT*)*×D*^.

#### Variance-weighted *R*^2^

We compute *R*^2^ per output dimension and then variance-weight across dimensions, i.e., the standard coefficient of determination with variance-weighted multioutput aggregation applied to (**Y**_flat_, **Ŷ** _flat_).

#### Correlation (Corr)

We compute Pearson correlation independently per dimension *d* ∈ {1, …, *D*} on the flattened vectors (**Y**_flat_[:, *d*], **Ŷ**_flat_[:, *d*]) and report the mean across dimensions. If either vector has near-zero standard deviation (*<* 10^−8^), we define the correlation for that dimension to be 0 to avoid numerical issues.

#### RMSE (in original kinematic units)

During training we may standardize targets using training-set statistics. For reporting RMSE in Tab. 2, we undo any target standardization and compute

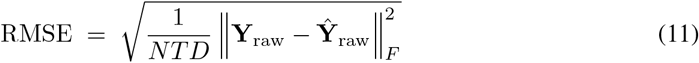

where ∥ · ∥_*F*_ denotes the Frobenius norm and **Y**_raw_, **Ŷ** _raw_ are in the original units.

#### Mean squared jerk (MSJ)

Given predictions 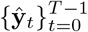 sampled at interval Δ*t*, we compute discrete jerk

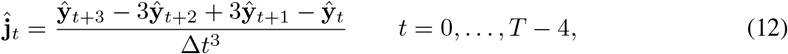

and

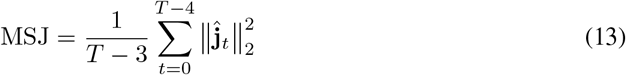

#### Aggregation across subjects/sessions

For datasets with multiple subjects/sessions, we follow the main-text protocol: compute metrics on each evaluation unit (subject/session) and then average to form a dataset-level number.

#### 95% confidence intervals over five seeds

For each metric, let *m*_1_, …, *m*_5_ be the five seed-level values. We report the mean 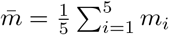 and a two-sided 95% t-interval:

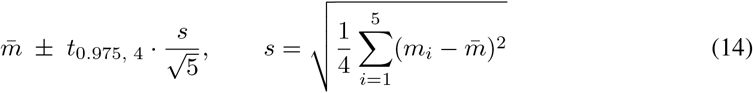

where *t*_0.975,4_ ≈ 2.776.

#### Error bars in Fig. 3b

We plot the mean across seeds with error bars equal to the standard error of the mean (s.e.m.) over five seeds:

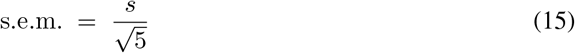

### B.3 Compute

All experiments were run on a single workstation:

- **CPU / RAM:** Intel Core i7-13700H; 32 GB RAM.
- **GPU:** NVIDIA RTX 4090.
- **Software:** Python 3.10; PyTorch 2.1; CUDA 12.1.
- **Runtime:** ∼2–6 hours per dataset for a single training run (depending on dataset size and model); prior sweeps and day-shift evaluation add ∼1–3 hours.

## C Windowing, Splits, and Input Construction

### C.1 Window Construction and Lag Indexing (Sec. 3.2)

All datasets are converted into supervised examples by sliding a causal window over each recording and extracting (i) a neural history window and (ii) a target kinematic segment aligned *within* the same history, exactly as defined in Sec. 3.2.

#### Seconds-to-steps conversion

For a recording with sampling interval Δ*t*, durations are converted to discrete steps by

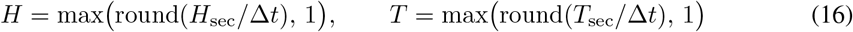

where *H*_sec_ is the history length in seconds and *T*_sec_ is the decoded horizon in seconds. In all main experiments we use within-window decoding and enforce *T < H*, so the decoding latency (*H* − *T*)Δ*t* is strictly positive. For the history-length sensitivity sweep in Appendix J.6, we additionally allow *T* ≤ *H* (including the zero-latency case *T* = *H*) purely for ablation; all main-text results and comparisons use the strict *T < H* setting.

#### Within-window targets and initialization

For each valid start index *t*_0_ (0-based), we construct

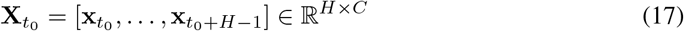

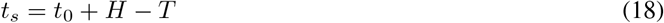

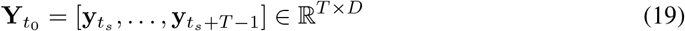

This corresponds to a fixed decoding latency of (*H* − *T*)Δ*t*. Consistent with Sec. 3.2, the SS-DMP rollout initial state is predicted from the causal neural window, rather than copied from the ground-truth target segment:

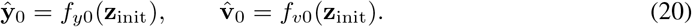

The corresponding supervised targets are the first position of the target segment,

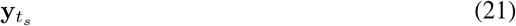

so that training encourages

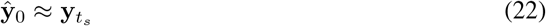

At inference time, no ground-truth kinematic state from the target segment is used.

#### Valid-window filtering (including masked intervals for spikes)

Some recordings provide a boolean validity mask **m** ∈ {0, 1} ^*n*^ indicating time indices with valid behavioral labels. For spiking benchmarks, this mask is derived from the dataset-provided evaluation mask and aligned to the binned time grid. We keep only window start indices *t*_0_ such that *all* timesteps required by the example are valid. In within-window mode, a sufficient condition is **m**[*t*_0_ : *t*_0_ + *H* − 1] = 1, which guarantees that both the neural history and the within-window target segment are fully supported.

#### Stride (non-overlapping decoded segments)

Start indices are subsampled by a stride *S* (in steps). In the main results, we use non-overlapping decoded segments for *all* datasets:

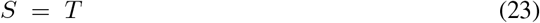

Thus successive decoded segments tile time without overlap, avoiding ambiguity in how window-level metrics relate to continuous performance.

#### Main protocol hyperparameters

All main results use (*H*_sec_, *T*_sec_) = (0.7 s, 0.5 s) (Sec. 4.2). Table 6 reports the resulting discrete step counts for the sampling intervals used in our benchmarks.

**Table 6:** Main protocol: seconds-to-steps conversion. All main experiments use (*H*_sec_, *T*_sec_) = (0.7 s, 0.5 s) and stride *S* = *T* .

| Dataset family | $\Delta t$ (s) | $H$ | $T$ | $S$ (steps) |
| --- | --- | --- | --- | --- |
| ECoG (MouseTrack, BCI IV-4) | 0.001 | 700 | 500 | 500 |
| Spikes (M2, H1) | 0.02 | 35 | 25 | 25 |

### C.2 Splits and Masks

#### ECoG splits (MouseTrack, BCI IV-4)

For each subject recording of length *n* samples (after alignment/resampling), we construct contiguous temporal splits by sample index: train {0, …, *n*_tr_ − 1}, validation {*n*_tr_, …, *n*_tr_ + *n*_va_ – 1}, and test {*n*_tr_ + *n*_va_, …, *n* – 1}, where *n*_tr_ = ⌊0.8*n*⌋ and *n*_va_ = ⌊0.1*n*⌋ (Sec. 4.2). Neural standardization (when enabled) is fit on the training portion and applied to validation/test within the same recording. MouseTrack metrics are averaged over four subjects; BCI IV-4 metrics are averaged over three subjects, consistent with Tab. 1.

#### Spike splits (M2, H1) and validation suffix

We follow the benchmark-defined session protocol: held-in calibration sessions are used for training/validation and held-out calibration sessions are used for testing (Sec. 4.2). Within each held-in session, we construct a contiguous train/validation split by time using a validation suffix (last 20% of samples), i.e., the first 80% of samples are used for training and the final 20% for validation.

#### Evaluation mask semantics and window validity

For spikes, the dataset-provided evaluation mask is aligned to the binned time grid and used as the recording validity mask. A window start *t*_0_ is considered valid iff every timestep required by the example is marked valid (Appendix. C.1); thus, all evaluated windows are fully supported by labeled kinematics under the within-window construction.

#### Day-shift evaluation protocol (M2; Sec. 4.4.3)

For the day-shift analysis, we evaluate each held-out session independently in chronological order, without weight updates and without using any held-out behavioral labels beyond what is required to compute evaluation metrics, matching the main-text protocol.

## D Datasets and Preprocessing (Extended)

### D.1 Dataset Details Beyond Tab. 1

This section provides *locked* dataset and preprocessing details sufficient to reproduce our inputs exactly from the public benchmark files, without relying on repository-specific implementation pointers.

#### MouseTrack (ECoG)[23]

MouseTrack is provided as four subject-specific `.mat` files (rh/gf/fp/rr), each containing an ECoG matrix **X** ∈ ℝ^*n×C*^ and continuous 2D cursor trajectories. In our release, channel counts are rh (64), gf (64), fp (61), rr (47). Neural signals are sampled at 1000 Hz; behavioral cursor trajectories are aligned by upsampling/interpolation to the same 1000 Hz grid, so the effective sampling interval is Δ*t* = 1 ms. We decode 2D cursor *position* in dataset units. When velocity is needed (e.g., SS-DMP initialization or prior fitting), it is derived consistently from the position trace as described in Appendix. D.5.

#### BCI Competition IV Dataset 4 (ECoG)[24]

BCI IV-4 is provided as three subject .mat files (sub1/sub2/sub3) containing labeled training data only. Each file provides ECoG time series **X** ∈ ℝ^*n×C*^ and 5D kinematic trajectories **Y** ∈ ℝ^*n×*5^; in our copy, channel counts are sub1 (62), sub2 (48), sub3 (64). Signals are nominally sampled at 1000 Hz; kinematic trajectories are aligned by upsampling/interpolation to the same 1000 Hz grid, yielding Δ*t* = 1 ms. Because the official competition test set is unlabeled, we evaluate only on labeled data with subject-wise temporal holdout (Sec. 4.2).

#### M2 (spikes)[25]

M2 is distributed as NWB files under held-in calibration sessions (for training/validation) and held-out calibration sessions (for testing). In the main setting we use held-in sessions for training/validation (7 sessions) and held-out sessions for testing (6 sessions). Each session contains spike times for a fixed set of 96 units and continuous 2D kinematics (*positions*) with timestamps in seconds. We use the benchmark-provided validity mask (eval_mask) to restrict evaluation to intervals with valid kinematic supervision (Appendix. C.2). Units for the kinematics are benchmark-defined arbitrary units (AU).

#### H1 (spikes)[26]

H1 is distributed as NWB files with 7D kinematic *positions* and a benchmark-provided validity mask. In the main setting we use held-in sessions for training/validation (13 sessions) and held-out sessions for testing (14 sessions). In our release, all sessions contain 176 units (the dataset supports up to ≤ 192 units in general). Kinematics are sampled at 50 Hz (i.e., Δ*t* = 0.02 s) as specified by NWB timing attributes; kinematic units are benchmark-defined arbitrary units. We decode the 7D kinematic *positions* (not velocity targets).

### D.2 ECoG Raw Stream

ECoG preprocessing is applied prior to windowing. Given raw ECoG 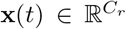 sampled at 1000 Hz, we apply:

- **Common-average referencing (CAR)**. We subtract the per-time-step channel mean: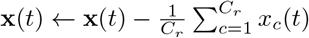
- **Bandpass filtering**. We used a filter to process the raw data, filtering it within the 1-200Hz range.
- **Notch filtering**. We apply an notch filter at 60 Hz with *Q*=30 (and harmonics up to Nyquist).
- **Sampling alignment**. For the ECoG benchmarks considered here, neural signals are sampled at 1000 Hz and behavioral trajectories are aligned by upsampling/interpolation to the same 1000 Hz grid. We therefore use a fixed Δ*t* = 1 ms for ECoG throughout.
- **Z-scoring**. For each ECoG subject recording, we fit feature mean and standard deviation on the training portion of the contiguous temporal split and apply the same normalization to validation and test (Appendix. C.2).

### D.3 PSD Stream and Alignment to Raw (Sec. 3.3)

When using raw+PSD fusion, we compute a PSD feature sequence over the *same* history window as the raw stream (after the preprocessing above). Given a history segment 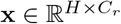 sampled at *f*_*s*_ = 1000 Hz, we construct PSD frames using Welch bandpower features:

- **Window/hop**. We use a Welch window length of 0.1 s and hop of 0.05 s, i.e., *n*_seg_ = round(0.1 *f*_*s*_) and hop *s* = round(0.05 *f*_*s*_). Frame start indices are 0, *s*, 2*s*, …, *H* − *n*_seg_ (at least one frame is produced).
- **Bands**. We use *B* = 25 non-overlapping bands spanning 1–100 Hz (4 Hz bins), yielding per-frame features in 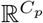 with *C*_*p*_ = *C*_*r*_ · *B* by concatenating band-power features across all channels.
- **Log and normalization**. We apply log(1 + ·) to band powers. When multiple PSD frames exist in a history window, we z-score each PSD feature dimension across frames *within the same window* (window-local; no behavioral labels). If a window contains only one PSD frame, we skip this window-local z-scoring. This PSD-specific normalization is separate from the global (training-set) z-scoring used for raw time-domain neural features, and we apply the same PSD pipeline for all models that use PSD inputs.

#### Alignment to the decoded segment start

For within-window decoding, the decoded segment starts at raw index *k*_*r*_ = *H* − *T* within the history (0-based; Sec. 3.2). To align the PSD sequence of length *H*_*p*_ to the same relative time, we map by normalized time:

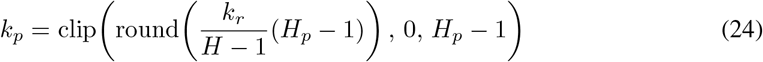

which matches the within-aligned query pooling defined in Sec. 3.3.

### D.4 Spike Preprocessing

Spike preprocessing is applied per session prior to windowing:

- **Binning**. Spike times are converted to counts on a regular grid with Δ*t*_bin_ = 20 ms by histogramming each unit’s spike times into bins. Kinematics and the benchmark validity mask are linearly interpolated to the same bin centers; the interpolated mask is thresholded at 0.5 to obtain a boolean validity mask (Appendix. C.2).
- **Causal exponential smoothing**. We apply causal exponential smoothing with time constant *τ*_*s*_ = 0.05 s:

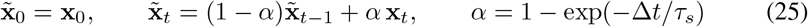

where Δ*t* = Δ*t*_bin_ is the post-binning timestep.
- **Z-scoring on held-in training data**. We compute a global mean and standard deviation from held-in *training* recordings (restricted to valid timesteps when the validity mask is present) and apply this normalization to held-in training and validation.
- **Label-free test-session re-normalization (day-shift analysis only)**. For the M2 day-shift analysis (Sec. 4.4.3), we additionally enable a label-free per-session re-normalization on held-out test sessions: mean and variance are recomputed from that session’s neural data only (restricted to valid timesteps) and used to re-standardize the session, without using behavioral labels. Unless explicitly stated, the main table results do not rely on this optional re-normalization.
- **No PSD for spikes**. We do not compute PSD features for spiking inputs.

### D.5 Target Preparation and Velocity Estimation

#### Target standardization and reporting units

We standardize kinematic targets using training-set mean and standard deviation and apply the same affine transform to validation/test targets. We store the training mean/std and undo this transform when reporting errors in original dataset units. (Note that *R*^2^ and Corr are invariant to applying the same affine transform to both predictions and targets; RMSE is reported after undoing standardization.) No additional scaling, clipping, or unit conversion is applied beyond this target standardization.

#### Target reconstruction

The pipeline include target reconstruction/smoothing step for visualization or downstream post-processing;

#### Savitzky–Golay smoothing

Given 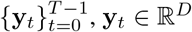, we apply a per-dimension Savitzky–Golay (SG) filter to obtain 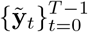. For each dimension, we fit a degree-*p* polynomial over a sliding window centered at *t* and replace *y*_*t,d*_ by the center value. We use *p* = 3 and a fixed temporal window *w*_sec_ = 0.1 s, implemented as an odd sample length *w* = 2 ⌊ (*w*_sec_*/*Δ*t*)*/*2⌋ + 1 (clipped at the segment length if needed). At sequence boundaries, we use standard SG boundary handling (interp/reflect-style) so the output length matches the input.

#### Implementation details

We apply SG independently to each dimension and each segment (no cross-segment leakage), and the filter is applied only to the target sequence used in this reconstruction module. When SG is used for derivative estimation in forcing inference (Stage 1), we apply it only to obtain stable finite-difference derivatives and never alter the definition of the goal **g** or the evaluation targets.

#### Velocity and acceleration

When velocity is required (e.g., SS-DMP initialization and prior fitting), we use a consistent one-step finite-difference operator on the position target:

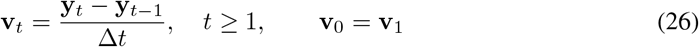

This convention ensures consistency between the target, SS-DMP initialization, and any behavior-derived prior computations that require derivatives.

## E Encoder Architecture Details

This section specifies the encoder used in Sec. 3.3 with *locked* architectural hyperparameters for exact reproducibility. The encoder uses a Conv–Transformer architecture applied independently to each stream (raw ECoG, PSD/band-power ECoG, or spikes), followed by within-aligned query pooling and, for ECoG, lightweight raw–PSD fusion.

### Main-setting hyperparameters (all datasets unless noted)

We use hidden size *h* = 256, GELU nonlinearity, dropout *p* = 0.1 throughout (input/conv/attention/FFN), *L*_conv_ = 2 temporal convolution blocks (kernel size *k* = 5, stride 1, padding to preserve length), and an *L*_tr_ = 4-layer Transformer encoder with *n*_head_ = 8 attention heads and feed-forward dimension *h*_ff_ = 4*h* = 1024. LayerNorm is applied in a pre-norm Transformer and as written below in the conv stack.

### E.1 Per-modality tokenization and projection

Index modalities by *m* ∈ {*r, p, s*}, where *r* denotes raw ECoG, *p* denotes PSD/band-power ECoG, and *s* denotes spikes. For a window start *t*_0_, each modality provides an input sequence 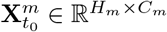, where *H*_*m*_ is the number of time steps/frames for modality *m* (note *H*_*r*_≠ *H*_*p*_≠ *H*_*s*_ in general). We form an initial token sequence by LayerNorm followed by a learned linear projection to ℝ^*h*^:

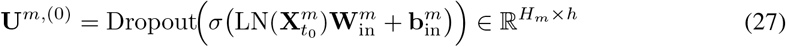

where *σ*(·) is GELU.

### E.2 Temporal convolution stack

We apply *L*_conv_ = 2 residual temporal conv blocks along the time axis (sequence length preserved). Each block uses a 1D convolution with kernel size *k* = 5, stride 1, and padding chosen to preserve length:

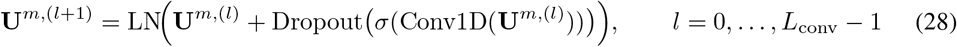

No dilation or downsampling is used in the main setting.

### E.3 Transformer encoder

After the conv stack, we apply a Transformer encoder with *L*_tr_ = 4 layers, *n*_head_ = 8 heads, and FFN dimension *d*_ff_ = 1024. We use sinusoidal positional encodings added to the token sequence. Let the Transformer input token sequence be

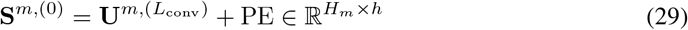

A single pre-norm Transformer layer *ℓ* ∈ {0, …, *L*_tr_ − 1} is:

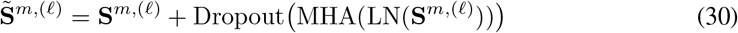

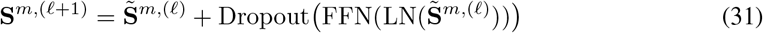

where MHA is multi-head self-attention and FFN is a two-layer MLP with hidden size *d*_ff_ and GELU activation. The encoded token sequence is the final-layer hidden states:

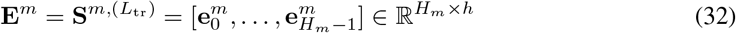

Self-attention is bidirectional *within* the observed window; the overall protocol remains window-causal because no neural samples beyond the end of the observed window are ever accessed.

### E.4 Within-aligned query pooling (Sec. 3.3)

In within-window decoding, the target segment starts at *t*_*s*_ = *t*_0_ + *H* − *T* (Sec. 3.2). Using 0-based indices within the raw history window of length *H*, the aligned raw index is

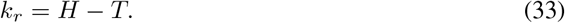

For modality *m* with sequence length *H*_*m*_, we map this alignment point by normalized time within the window:

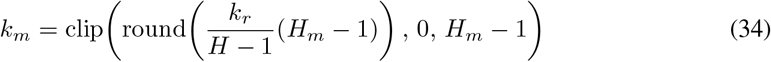

We define the initialization embedding as the aligned token,

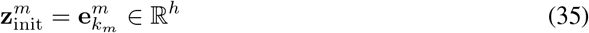

and obtain a global control embedding by using 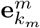 as a length-1 query attending over the full encoded sequence:

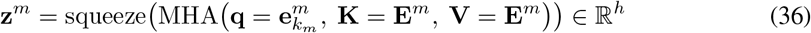

This pooling explicitly anchors both 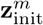 and 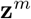 to the decoded segment start, matching the SS-DMP initialization at *t*_*s*_ (Sec. 3.2).

### E.5 Multimodal fusion for ECoG (raw+PSD)

For ECoG fusion, we encode raw and PSD streams independently and obtain 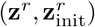 and 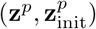. We fuse them with lightweight MLPs:

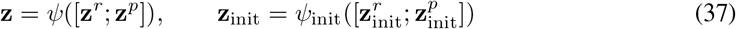

In the main setting, both *ψ* and *ψ*_init_ are two-layer MLPs:

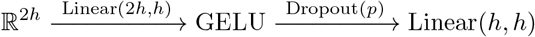

with *p* = 0.1. For spiking data, we use a single stream and set (**z, z**_init_) = (**z**^*s*^, **z**^*s*^).

## F Supplement to the SS-DMP Generator

This section provides the implementation-exact details for the SS-DMP generator used in Sec. 3.5. We emphasize **main-text consistency**: (i) the rollout uses the **forward-Euler** discretization for the transformation system (Eq. 5); (ii) the phase update uses the **closed-form exponential** update; (iii) time scale is fixed to *τ* ≡ 1 in all main experiments (physical time expressed via Δ*t*).

### F.1 RBF Basis Construction (Centers and Widths)

Let *K* denote the number of basis functions and let the rollout horizon be *T* discrete steps with sampling interval Δ*t*. The *i*-th Gaussian RBF is

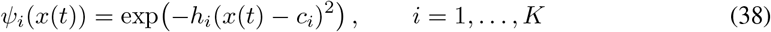

#### Time scale *τ*

All main results use *τ* ≡ 1 (fixed across datasets). Thus any appearance of *τ* in the DMP parameters is interpreted under *τ* = 1 and physical time is carried by Δ*t*.

#### Centers *c*_*i*_ (time-uniform, mapped to phase)

We place *K* centers uniformly over the *discrete* segment time range {0, …, *T* – 1} and map each time to phase using the canonical decay (*τ* = 1). Define

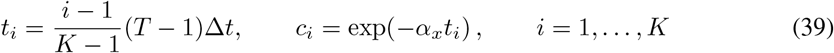

which matches Sec. 3.5 exactly.

#### Widths *h*_*i*_ (overlap in phase domain)

We set widths using adjacent center spacing in the *phase* domain to ensure smooth overlap:

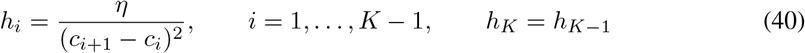

where *η* is a fixed overlap constant (main setting: *η* = 1). This is the width schedule used by the SS-DMP generator described in Sec. 3.5.

#### Normalized RBF mixture

Given weights **w** ∈ ℝ^*D×K*^, the generator forms the normalized mixture

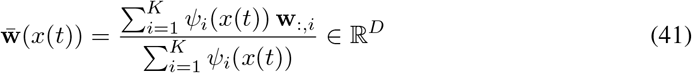

### F.2 Phase Update, Forcing, and Euler Rollout

#### Per-dimension independence

The generator applies the same second-order dynamics *independently* to each output dimension *d* ∈ {1, …, *D*}. Equivalently, the full system over **s**_*t*_ ∈ ℝ^2*D*^ is block-diagonal with *D* uncoupled 2 × 2 blocks; there are no cross-dimension coupling terms in the generator matrices.

#### Phase update

The scalar phase is initialized at *x*_0_ = 1 and updated by

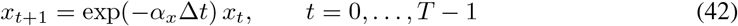

(Recall *τ* = 1 in the main setting.)

#### Forcing input (exactly as in Sec. 3.5)

Given weights **w** ∈ ℝ^*D×K*^, goal **g** ∈ ℝ^*D*^, and initial position **y**_0_ ∈ ℝ^*D*^, the forcing is

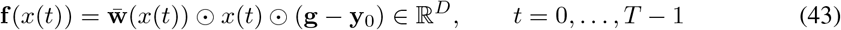

where 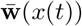 is defined in Eq. (41).

#### Continuous-time base matrices specialized to *τ* = 1

For each dimension *d*, define state **s**_*d*_(*t*) = [*y*_*d*_(*t*); *v*_*d*_(*t*)] with 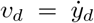 and input *u*_*d*_(*t*) = [*g*_*d*_; *f*_*d*_(*x*(*t*))]. The continuous-time SS-DMP dynamics are

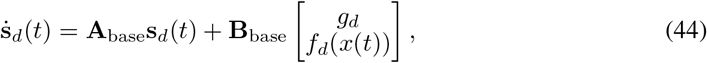

with

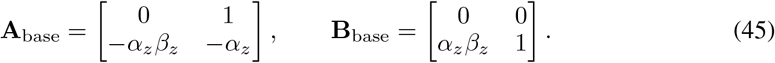

These are the base SS-DMP matrices from Sec. 3.5 specialized to *τ* = 1.

#### Euler discretization and output indexing (main-text semantics)

Let the discrete state be **s**_*d,t*_ = [*y*_*d,t*_; *v*_*d,t*_] with initialization **s**_*d*,0_ = [*y*_0,*d*_; *v*_0,*d*_]. Each step applies forward Euler with step size Δ*t*:

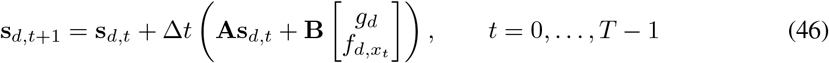

The generator returns the *length-T segment*

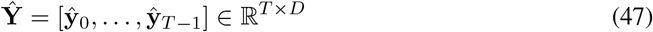

where **ŷ**_*t*_ is the position component of **s**_*t*_. This matches Sec. 3.2, where **Ŷ**_*t*_ is aligned to the target sample 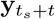

#### Within-window initialization alignment (Sec. 3.2)

In within-window decoding, the target segment starts at *t*_*s*_ = *t*_0_ + *H* − *T* . The generator is initialized at the segment start:

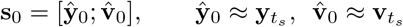

and the returned 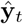 corresponds to the target 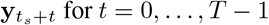 (as stated in Sec. 3.2).

#### Subject-specific dynamics injection (main-text blending with separate *λ*_*A*_, *λ*_*B*_)

When behavior priors are enabled (Sec. 3.6), the generator uses per-dimension matrices **A**_(*s,d*)_ ∈ ℝ^2*×*2^ and **B**_(*s,d*)_ ∈ ℝ^2*×*2^ and blends them with the base dynamics using the fixed main-text coefficients:

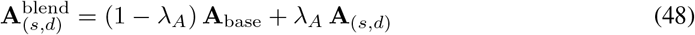

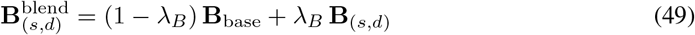

The Euler step in Eq. (46) is then applied with 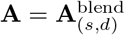 and 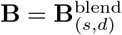 independently for each dimension *d*. In the main setting, we use fixed defaults *λ*_*A*_ = 0.2 and *λ*_*B*_ = 0.5 unless explicitly stated otherwise.

#### Dataset-dependent Δ*t*

The generator step size Δ*t* equals the timestep of the input/target series after dataset-specific alignment/preprocessing (Appendix. D): ECoG uses Δ*t* ≈ 1 ms after alignment, while spiking datasets use the post-binning timestep (main setting: Δ*t* = 0.02 s).

## G Supplement to Behavior-Derived Subject Priors

This appendix provides a complete, reproducible description of the behavior-derived subject prior fitting procedure. The goal is to estimate subject-specific continuous-time dynamics matrices 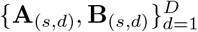 *using training kinematics only*, consistent with the SS-DMP generator family in Sec. 3.5. All fitting described below is performed separately for each subject *s* and uses only windows drawn from that subject’s *training split* (no validation/test behavior is used at any stage).

### G.1 Stage 1: From 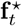 to generator-consistent w^⋆^

Stage 1 infers a forcing representation that is *consistent with the SS-DMP forcing family* (Eq. (3)), so that the downstream prior fitting in Stage 2 is aligned with the same forcing parameterization encountered at test time.

#### Within-window segment and goal definition

Consider one training target segment of length *T* (within-window decoding; Sec. 3.2)

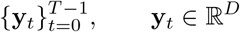

sampled at interval Δ*t*. We set the goal to the segment endpoint

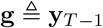

To obtain stable derivative estimates, we use the next sample **y**_*T*_ (immediately after the segment) *only* for finite differences; **y**_*T*_ is never used as a learning target.

#### Smoothing and finite differences (training-behavior only)

To stabilize behavioral derivatives, we smooth positions per dimension with a Savitzky–Golay filter (order *p*=3, fixed temporal window *w*_sec_=0.1 s implemented as an odd sample length *w* per dataset), yielding 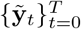. We then compute velocities and accelerations by finite differences:

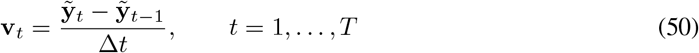

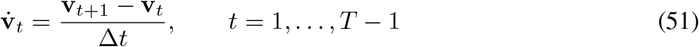

All quantities above use only training kinematics.

#### Continuous-time forcing target 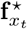

Rearranging the canonical DMP transformation system (Eq. (2)) yields the forcing target (on steps *t* = 1, …, *T* − 1):

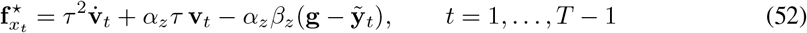

#### SS-DMP forcing family and linear design matrix

The SS-DMP restricts forcing to the RBF family induced by Eq. (3). Let {*x*_*t*_} be the phase sequence generated by Eq. (42) with *x*_0_ = 1. Let 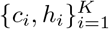 be the basis centers/widths defined exactly as in the generator (Sec. 3.5; see also Appendix F.1 for the fixed construction). Define activations and their normalized versions:

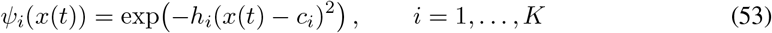

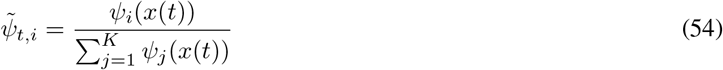

For each step, define the feature row

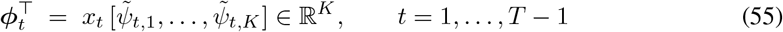

and stack them to obtain **Φ** ∈ ℝ^(*T* −1)*×K*^ .

Under Eq. (3), letting **y**_0_ denote the segment start (here 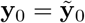), the generator-consistent forcing for dimension *d* is linear in its basis weights:

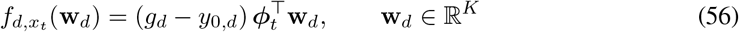

#### Ridge projection to obtain w^⋆^ (per window, per dimension)

For each window and each output dimension *d* ∈ {1, …, *D*}, define the displacement scale *s*_*d*_ ≜ *g*_*d*_ − *y*_0,*d*_. We fit 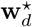 by ridge regression over steps *t* = 1, …, *T* − 1:

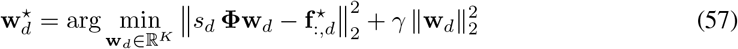

with *γ* = 10^−3^ (fixed across all main experiments, matching the main text). When *s*_*d*_ *<* 10^−6^ (near-zero displacement), we set 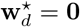 for numerical stability. We then define the generator-consistent forcing used downstream by

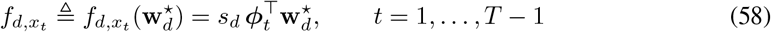

#### Why this projection matters

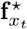 in Eq. (52) is an *unconstrained* forcing implied by the continuous-time DMP equation. However, the SS-DMP generator can only realize forcings of the form in Eq. (3). Stage 1 therefore projects 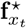 onto the realizable RBF family, ensuring Stage 2 learns subject dynamics using the same forcing parameterization that will be produced at test time from predicted **ŵ** .

### G.2 Stage 2: Constrained ridge regression for (A_(*s,d*)_, B_(*s,d*)_)

Stage 2 fits continuous-time dynamics independently for each output dimension *d* and each subject *s* by stacking samples from *all valid training windows* of that subject (not session-specific fitting).

#### Regression model (per dimension)

Using tuples 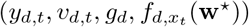 and 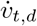 from Stage 1 (for *t* = 1, …, *T* − 1), we fit:

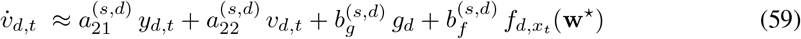

Stacking all samples across all training windows yields a ridge system **X**_*d*_ ∈ ℝ^*n×*4^, **y**_*d*_ ∈ ℝ^*n*^ with rows 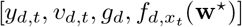 and targets 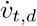.

#### Objective and stability constraints (continuous time)

We solve a *constrained* ridge regression:

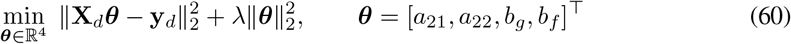

subject to the continuous-time stability conditions for the second-order companion form

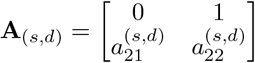

The main text enforces tr(**A**_(*s,d*)_) *<* 0 and det(**A**_(*s,d*)_) *>* 0. For this structured 2 × 2 form, these are equivalent to the simple sign constraints

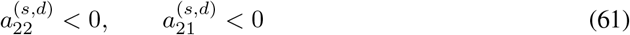

since tr(**A**) = *a*_22_ and det(**A**) = −*a*_21_. We implement the strict inequalities as *a*_21_ ≤ −*ϵ*_stab_ and *a*_22_ ≤ −*ϵ*_stab_ with *ϵ*_stab_ = 10^−6^.

We use the same ridge penalty as in Stage 1, *λ* = 10^−3^, matching the main-text setting. Since we roll out with a small step size Δ*t*, the corresponding discrete-time update typically preserves stability in practice for this model.

#### Exact solution procedure (small QP with two active constraints)

Eq. (60) with constraints (61) is a strictly convex quadratic program. Because there are only two inequality constraints (on *a*_21_ and *a*_22_), it can be solved exactly by checking a small number of active-set cases:

1. Compute the unconstrained ridge solution 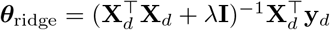. If it satisfies *a*_21_ ≤ −*ϵ*_stab_ and *a*_22_ ≤ −*ϵ*_stab_, accept it.
2. Otherwise, solve the constrained optimum by considering which of the two constraints are active: (a) fix *a*_21_ = − *ϵ*_stab_ and optimize the remaining variables; (b) fix *a*_22_ = − *ϵ*_stab_ and optimize the remaining variables; (c) fix both *a*_21_ = *a*_22_ = − *ϵ*_stab_ and optimize (*b*_*g*_, *b*_*f*_). Each case reduces to an unconstrained ridge problem in the remaining free variables (a small linear system).
3. Among feasible candidates, choose the one with the smallest objective value in Eq. (60). This yields the exact minimizer under the stated constraints.

#### Assembling matrices

For each dimension *d*, we assemble the fitted continuous-time blocks:

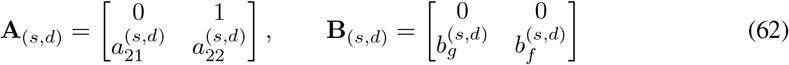

We store **A**_(*s*)_ ∈ ℝ^*D×*2*×*2^ and **B**_(*s*)_ ∈ ℝ^*D×*2*×*2^ as the subject prior.

#### Learning modes for ablations

To support the ablations in Tab. 3, we consider: (i) **A***-only* fitting: constrain and fit (*a*_21_, *a*_22_) while setting (*b*_*g*_, *b*_*f*_) to the base DMP values; (ii) **B***-only* fitting: fit (*b*_*g*_, *b*_*f*_) while setting (*a*_21_, *a*_22_) to the base DMP values; (iii) **A** + **B** (main setting): fit all four coefficients as above. In all cases, when **A** is learned we enforce the continuous-time stability constraints.

### G.3 Using the prior in the generator (fixed *λ*_*A*_, *λ*_*B*_)

During decoding, we inject the learned prior into the SS-DMP generator by fixed blending, exactly as in Eq. (10):

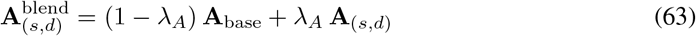

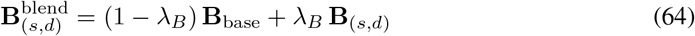

Unless stated otherwise, we use fixed defaults (*λ*_*A*_, *λ*_*B*_) = (0.2, 0.5) and do not perform any test-time fitting, calibration, or access to held-out behavioral labels.

### G.4 Sanity checks and common failure modes

#### Stage 1 reconstruction sanity check

A practical sanity check is that rolling out the SS-DMP using training-derived 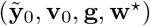 reconstructs the observed segment 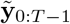 with low RMSE (up to discretization and derivative noise). This detects ill-conditioned cases where the fixed RBF family cannot represent the forcing shape.

#### Common failure modes

Two frequent issues are: (i) near-zero displacement 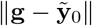 (handled by setting **w**^⋆^ = **0** for those dimensions); (ii) noisy accelerations from finite differences (mitigated by Savitzky–Golay smoothing and ridge regularization). Because all fitting uses only training kinematics, these checks and mitigations do not introduce evaluation leakage.

## H Loss and Optimization (Full Details)

### H.1 Training Objective

This section *locks* the exact training objective used for all main-text results.

#### Notation

Let **Ŷ**, **Y** ∈ ℝ^*B×T ×D*^ denote the predicted and ground-truth kinematic segments for batch size *B*, horizon length *T* (discrete steps), and output dimension *D*. All losses average over *all* tensor entries.

#### Position MSE (only loss used)

In the main setting, we train *only* with segment-wise position mean-squared error:

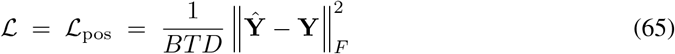

No auxiliary loss terms (endpoint/startpoint, velocity, smoothness, or parameter supervision) are enabled in the reported experiments. Model selection uses the lowest validation ℒ_pos_.

### H.2 Optimization and Model Selection (Main Setting)

This section fixes the optimization procedure corresponding to the main-text protocol.

#### Optimizer

We optimize all trainable parameters with AdamW:

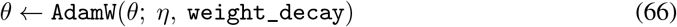

using learning rate *η* = 3 × 10^−4^ and weight decay 10^−2^. All other AdamW hyperparameters use PyTorch defaults (betas (0.9, 0.999) and *ϵ* = 10^−8^).

#### Learning-rate schedule (cosine with warmup)

We use a cosine decay schedule with linear warmup. Let *N* be the total number of optimizer updates in a run (i.e., the number of minibatches across all epochs), and let *N*_warm_ = ⌈0.05 *N*⌉ (5% warmup). For update index *u* ∈ {1, …, *N*}, the learning rate is

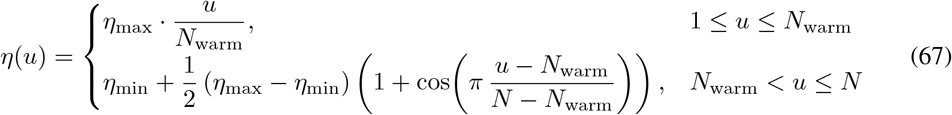

with *η*_max_ = 3 × 10^−4^ and *η*_min_ = 0. The scheduler steps *per update* (not per epoch).

#### Batching, epochs, and early stopping

We train with batch size 64 for up to 200 epochs. We apply early stopping on validation ℒ_pos_ with patience 20 epochs, and we select the checkpoint with the lowest validation loss for final test evaluation.

## I Baselines and Fairness (Implementation Details)

This appendix specifies the baselines used in our main comparisons (Sec. 4.3; Tab. 2) and the procedure used to ensure a fair, apples-to-apples evaluation.

### Fairness protocol (shared across all methods)

Unless explicitly stated otherwise, every baseline: (i) uses the *same* within-window windowing protocol, splits, masks, and input construction as Neural SS-DMP (Appendix C–C.2); (ii) predicts the same target trajectory segment **Y** ∈ ℝ^*T ×D*^ at the same horizon length *T* and sampling interval Δ*t*; (iii) uses the same training/validation selection rule and is evaluated with the identical metrics and uncertainty reporting (Appendix B.2); and (iv) shares the same seed protocol as the main text (same number of runs and identical data splits across seeds).

### Training objective for direct-trajectory baselines

All neural trajectory-regression baselines (Transformer/TCN/RNN/LSTM[5, 27, 6, 7]) are trained with the same trajectory loss configuration as the main setting (Appendix H). In particular, we do *not* use any auxiliary supervision on DMP parameters (no (**w, g**) targets) for these baselines.

### Which baselines are included

We report: (i) a linear Gaussian Kalman filter (KF); (ii) deep direct trajectory regressors (Transformer, TCN, RNN/LSTM); (iii) NoMAD[8], a drift-robust latent-dynamics alignment decoder for multi-session spiking datasets; and (iv) Regress+Project, a post-hoc projection baseline used in Tab. 3.

### I.1 Kalman Filter (KF)

We include a classic linear-Gaussian state-space decoder as a non-neural baseline.

#### State, observation, and model

We use a constant-velocity kinematic state **s**_*t*_ = [**y**_*t*_; **v**_*t*_] ∈ ℝ^2*D*^ and treat neural features **x**_*t*_ ∈ ℝ^*C*^ as observations:

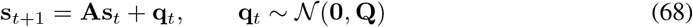

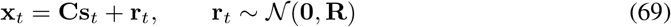

#### Fitting A and C (ridge-regularized least squares)

From the training split only, we build supervised pairs (**s**_*t*_, **s**_*t*+1_, **x**_*t*_). We estimate **A** and **C** by ridge regression:

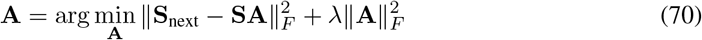

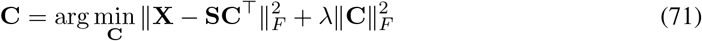

where rows of **S** stack **s**_*t*_, rows of **S**_next_ stack **s**_*t*+1_, and rows of **X** stack **x**_*t*_. We then estimate **Q** and **R** from residual covariances (training split only) with a small diagonal jitter for numerical stability.

#### Decoding under within-window evaluation

Given a within-window evaluation example with history 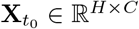, we run the Kalman filter forward over the *H* neural steps and take the last *T* filtered position means as the decoded segment (aligned to the within-window target definition in Appendix C.1). We initialize the filter with mean **0** and covariance 10^3^**I** (a diffuse prior).

### I.2 Neural Trajectory Regression Baselines

These baselines map the same neural history window to a *T* -step kinematic segment directly:

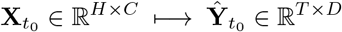

and do not use the SS-DMP generator.

#### Shared decoding head (time-query cross-attention)

To standardize the output interface across neural encoders, we use a shared cross-attention head. We create a normalized time grid *u*_1_, …, *u*_*T*_ ∈ [0, 1], embed it with a small MLP to form queries, and cross-attend to an encoder-produced memory sequence **M**:

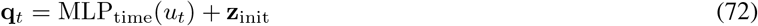

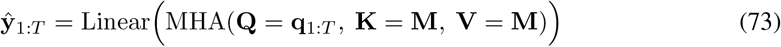

The initialization embedding **z**_init_ is taken from the within-aligned index (the same normalized-time mapping used for within-window alignment in Sec. 3.3 and Appendix D.3).

#### Transformer baseline

A pre-norm Transformer encoder (no causal mask) processes the raw stream (and, for ECoG settings that include PSD, an additional PSD stream; see below), followed by the shared cross-attention head.

#### TCN baseline

A causal dilated temporal convolutional network (TCN) encoder with exponentially increasing dilations encodes the history window, followed by the shared cross-attention head. Left-padding enforces causality.

#### Recurrent baselines (RNN/LSTM)

An RNN-family encoder (vanilla RNN or LSTM) processes the history window and outputs a memory sequence that is consumed by the same cross-attention head.

#### Raw+PSD handling for ECoG

When the main setting provides both raw and PSD inputs, baselines that support multimodal inputs encode the two streams separately and fuse them at the representation level before decoding. This ensures they have access to the same information content as Neural SS-DMP under the corresponding input setting.

#### Optimization and selection

All neural baselines follow the main-text optimization protocol (Appendix H.2): same optimizer family, epoch budget, batch size, gradient clipping, and validation-based checkpoint selection. Hyperparameters are fixed according to the main-text baseline configuration (i.e., not tuned per dataset unless explicitly stated in the main text).

### I.3 NoMAD (Spiking Drift-Robust Baseline)

We include NoMAD [8] only for *multi-session spiking* datasets, where distribution shift across sessions (“day shift”) is a primary challenge. NoMAD is a latent-dynamics alignment approach: it learns a *reference* dynamical latent model from held-in training sessions, then adapts to a held-out test session using *unlabeled* neural activity by aligning the test-session spiking distribution into the same latent dynamical space.

#### High-level procedure

NoMAD comprises two phases:

- **Reference training on held-in sessions (labeled)**. We fit a VAE-style dynamical latent model (LFADS-based in the released NoMAD implementation) on the held-in *training split* spanning multiple sessions/days, using spiking observations paired with kinematics for training and hyperparameter selection (within held-in data only). We then train a behavioral decoder from the inferred latent states (e.g., generator/factor states) to kinematics using the same held-in labeled data. The held-out test session is *never* used in this phase.
- **Held-out test-session alignment (unlabeled)**. For a held-out test session, we keep the reference dynamical model and the held-in behavioral decoder *fixed*, and update only a session-specific alignment/read-in module that maps test-session spiking activity into the reference latent space. This alignment uses *neural activity only* (no test-session behavioral labels) by optimizing unsupervised objectives on spiking data (restricted to valid intervals when an evaluation mask is present), e.g., maximizing spiking likelihood under the frozen dynamical model and matching latent distributions to the reference space as in NoMAD [8]. After alignment, the fixed held-in behavioral decoder is applied to the aligned test-session latents to produce kinematics.

#### Fairness and data-use constraints

Consistent with our leakage-prevention policy (Appendix B.1), NoMAD uses *no* behavioral labels from held-out test sessions for alignment, model selection, or early stopping. All hyperparameters are set using held-in data only, and test-session alignment is run with a fixed preset schedule (no tuning on test-session kinematics).

#### Scope in our experiments

We report NoMAD as a specialized drift-robust baseline on multi-session spiking benchmarks alongside KF and neural direct regressors. For ECoG datasets with single-recording temporal splits, we do not include NoMAD.

### I.4 Transformer+EMA: Causal Post-hoc Smoothing Control

Transformer+EMA is a diagnostic post-hoc smoothing baseline used in Table 3. It tests whether the gains of Neural SS-DMP can be explained by ordinary temporal smoothing of a direct Transformer regressor. The baseline first trains the direct Transformer under the same within-window protocol, data splits, preprocessing, optimizer, early stopping rule, and validation-based checkpoint selection as the Transformer baseline in Table 2. After the Transformer produces a trajectory prediction, we apply a causal exponential moving average (EMA) smoother to the predicted kinematic sequence. No model weights are updated during this smoothing step.

Let

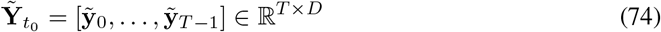

denote the trajectory predicted by the direct Transformer for one evaluation window. Transformer+EMA computes a smoothed trajectory

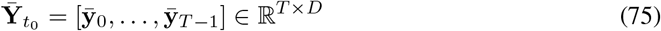

by applying the following recursion independently to each output dimension:

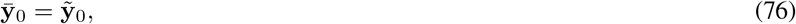

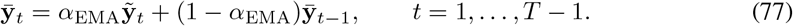

Here *α*_EMA_ ∈ (0, 1] controls the amount of smoothing: smaller values produce stronger temporal smoothing, while *α*_EMA_ = 1 recovers the original Transformer prediction.

To make the smoothing strength comparable across datasets with different sampling intervals, we parameterize the EMA by a physical time constant *τ*_EMA_. For a dataset with sampling interval Δ*t*, the EMA coefficient is

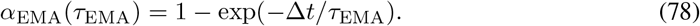

For each dataset and random seed, *τ*_EMA_ is selected on the validation split from the pre-specified grid

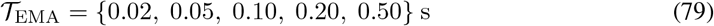

by minimizing validation position MSE after smoothing. The selected value is then fixed before test evaluation. Test-set behavioral labels, test-set smoothness statistics, and ground-truth test kinematics are never used to choose *τ*_EMA_ or *α*_EMA_.

The EMA smoother is causal with respect to the decoded trajectory index: the smoothed value 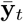 depends only on Transformer predictions 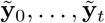 within the decoded segment. It does not use future ground-truth kinematics, future neural samples beyond the observed input window, or any test-session behavioral labels. When target variables are affinely standardized during training, applying Eq. (77) before or after undoing the standardization is equivalent, because the EMA recursion is affine and its weights sum to one at every step.

This baseline is intentionally simple and conservative. It isolates the effect of causal low-pass smoothing applied to an already trained direct regressor. Therefore, if Transformer+EMA reduces mean squared jerk but does not recover the accuracy of Neural SS-DMP, the remaining gap cannot be attributed to ordinary post-hoc smoothing alone.

### I.5 Regress+Project (Tab. 3)

Regress+Project tests whether Neural SS-DMP gains could be explained purely by projecting a regressed trajectory onto the SS-DMP trajectory family.

#### Step 1: direct trajectory regression

Train a direct neural regressor (the same Transformer baseline above) to predict a kinematic segment 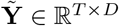 from the neural history window.

#### Step 2: fit generator parameters to 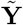

For each window, set the goal to the predicted end-point 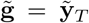 and use the same within-window initialization (**y**_0_, **v**_0_) as in the main protocol (Appendix C.1). Then infer 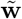 by the same generator-consistent ridge projection used elsewhere in the paper (Appendix G.1): we fit 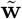 so that the induced forcing family reproduces 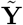 as closely as possible in least squares.

#### Step 3: re-rollout with the SS-DMP generator

Roll out the SS-DMP generator using 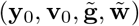 and the same discretization conventions as the main model (Appendix F.2), producing the projected trajectory **Ŷ** ^proj^.

#### Interpretation

If improvements were solely due to enforcing the SS-DMP constraint as a post-hoc projection operator, then Regress+Project would match Neural SS-DMP. A consistent gap supports the benefit of learning in the structured control space rather than projecting after unconstrained regression.

## J Extended Results (Supplementary)

### J.1 Additional Visualization Results

To complement the aggregate metrics in Tab. 2, we show additional time-series visualizations on all four datasets using the same evaluation protocol as the main table (HI → HO for spikes; held-out test split for ECoG). Fig. 4 summarizes one representative within-window decoded segment for each dataset (M2, H1, BCI IV-4, MouseTrack; Figs. 4a–4d), plotting three curves per dimension: ground truth, Neural SS-DMP, and the baseline.

**Figure 4:**
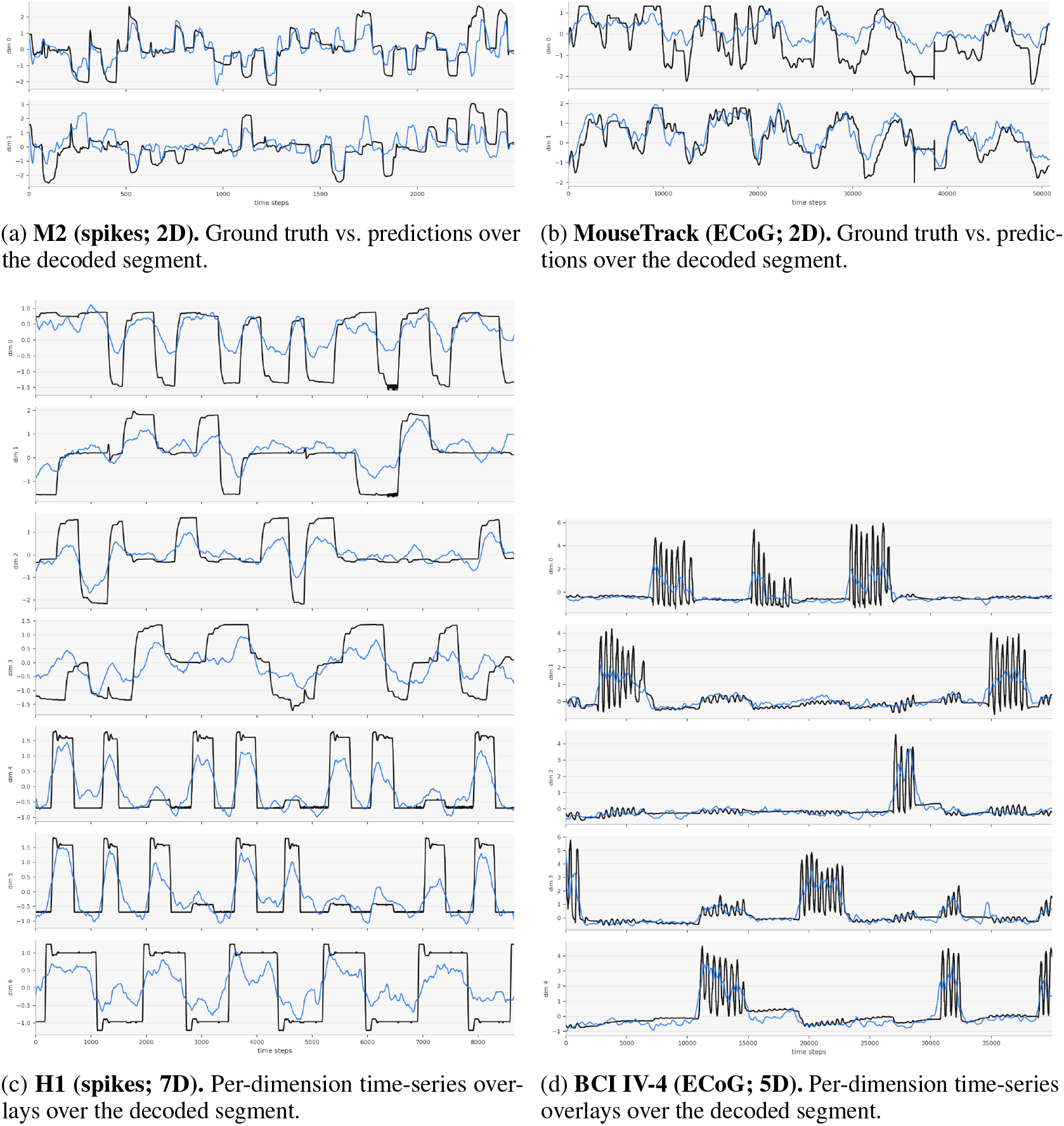
Visualization results across datasets. Each panel shows a representative decoded segment (median-*R*^2^ window under Neural SS-DMP on the corresponding split). Plots are reported in original kinematic units.

#### Selection protocol (non-cherry-picked)

For each dataset, we fix a random seed and choose a single representative test window whose Neural SS-DMP variance-weighted *R*^2^ is closest to the median across all test windows on the corresponding split. All curves are plotted in original kinematic units after undoing any target standardization.

#### What the curves show

Across datasets, Neural SS-DMP closely follows the ground-truth segment over the decoded horizon under the within-window protocol. In the 2D tasks (M2 and MouseTrack; Figs. 3a and 4b), the predicted trajectories match the overall path while avoiding sudden spikes or jagged transients. In the higher-dimensional tasks (BCI IV-4 and H1; Figs. 4d and 4c), the per-dimension overlays track the corresponding channels without large phase shifts or channel-specific instabilities, indicating consistent multi-DoF decoding under the same protocol.

#### Takeaway

These visualizations provide an intuitive sanity check that improvements in Tab. 2 correspond to meaningful trajectory fidelity over time, rather than being artifacts of aggregate statistics.

### J.2 Planned paired permutation tests for main *R*^2^ comparisons

To supplement the aggregate results in Table 2, we report planned paired permutation tests on variance-weighted *R*^2^. These tests are used as supportive evidence for the main accuracy comparisons under the reported five-seed evaluation protocol, rather than as an omnibus claim or definitive population-level inference that Neural SS-DMP is significantly better than every baseline on every dataset.

For each planned comparison between Neural SS-DMP and a baseline, we compute paired differences

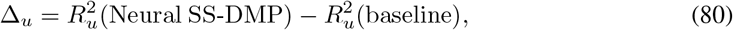

where *u* indexes a matched seed–evaluation-unit pair evaluated under the same split, preprocessing, and non-overlapping target windows. For ECoG datasets, the evaluation unit is a subject, and each subject is evaluated under five matched random seeds. For spiking datasets, the evaluation unit is a held-out session or held-out date, again evaluated under matched random seeds. Because random seeds share the same underlying evaluation units, these tests should be interpreted as supportive paired analyses of the reported experimental protocol rather than definitive population-level inference across independent subjects or sessions. The test statistic is the mean paired difference,

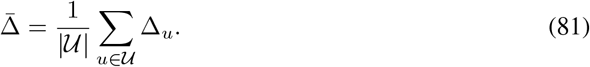

Under the null hypothesis of no systematic model advantage, we form a sign-flip permutation distribution by multiplying each Δ_*u*_ by an independently sampled sign in {−1, +1} and recomputing 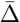. We use two-sided tests with the permutation *p*-value

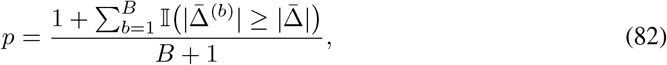

where 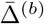 is the statistic after the *b*-th sign flip. We use *B* = 10,000 sign-flip samples with a fixed random seed. The reported *p*-values are uncorrected because the comparisons are pre-specified and used as supportive analyses.

We pre-specify two sets of comparisons. First, we compare Neural SS-DMP with the direct Transformer baseline on all datasets, because this isolates the effect of replacing unconstrained trajectory regression with the SS-DMP output generator while keeping the input features, training budget, and validation protocol matched. Second, on spiking datasets, we compare Neural SS-DMP with NoMAD as a specialized cross-session alignment baseline. Table 7 reports the resulting two-sided *p*-values.

**Table 7:** Planned paired permutation tests on variance-weighted *R*^2^. We report two-sided paired sign-flip permutation tests for the main planned accuracy comparisons. The tests compare Neural SS-DMP against the direct Transformer baseline on all datasets and against NoMAD on spiking datasets. *p*-values are uncorrected because these are pre-specified supportive comparisons.

| Dataset | Comparison | Two-sided $p$-value |
| --- | --- | --- |
| M2 | Neural SS-DMP vs. Transformer | 0.008 |
| MouseTrack | Neural SS-DMP vs. Transformer | 0.021 |
| H1 | Neural SS-DMP vs. Transformer | 0.071 |
| BCI IV-4 | Neural SS-DMP vs. Transformer | 0.18 |
| M2 | Neural SS-DMP vs. NoMAD | 0.021 |
| H1 | Neural SS-DMP vs. NoMAD | 0.23 |

The planned tests support the clearest *R*^2^ gains on M2 and MouseTrack, while indicating that the H1 and BCI IV-4 accuracy differences should be interpreted more cautiously. Together with Table 2, these tests support the conclusion that Neural SS-DMP improves the accuracy–smoothness trade-off, with statistically supported *R*^2^ gains in the strongest planned comparisons and competitive performance in the remaining settings.

### J.3 Controlled mechanistic validation of DMP trajectory generation

This appendix provides a controlled, decoder-independent validation of the DMP generator used in Neural SS-DMP. The purpose is not to claim that DMP is the only or universally optimal movement model, but to verify three properties that are desirable for an output-side movement generator: (i) the primitive variables can modulate both trajectory shape and endpoint, (ii) the stable dynamics recover from state perturbations without explicit replanning, and (iii) the generated motion exhibits smooth point-to-point kinematics with an accelerate–high-speed–decelerate temporal profile. This analysis is independent of neural decoding and therefore isolates the behavior of the DMP generator itself.

#### Setup

All experiments use a 2D point-to-point movement from start (0, 0) over a duration of *T* = 1.0 s with 301 time steps. For each coordinate, the DMP rollout follows the second-order transformation system used by our SS-DMP generator:

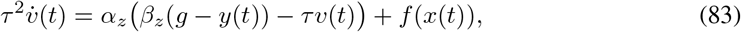

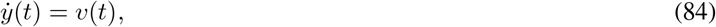

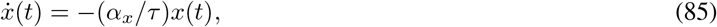

where *g* is the goal and *x*(*t*) is the canonical phase variable. The forcing term is represented by normalized radial basis functions,

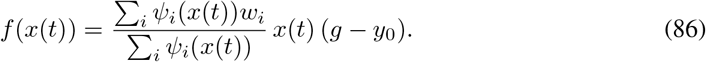

Thus, *g* controls the attracting endpoint, while *w* controls the smooth shape modulation around the attractor. We use *K* = 10 Gaussian basis functions unless otherwise stated.

#### Shape modulation by *w*

To verify that the forcing weights control path geometry, we fixed the start and goal to (0, 0) and (1, 1) and generated multiple DMP rollouts with different fitted forcing weights. As shown in Fig. 5a, changing *w* produces straight, curved, and S-shaped movements while keeping the same endpoint. Across the five trajectories, the mean terminal error was 7.45 × 10^−4^ and the maximum terminal error was 1.04 × 10^−3^. This confirms that the forcing term expands the generator beyond a rigid attractor while preserving goal convergence. We emphasize that this is an expressivity sanity check within the chosen RBF basis family, not a formal claim that a finite-basis DMP can represent every possible curve exactly.

**Figure 5:**
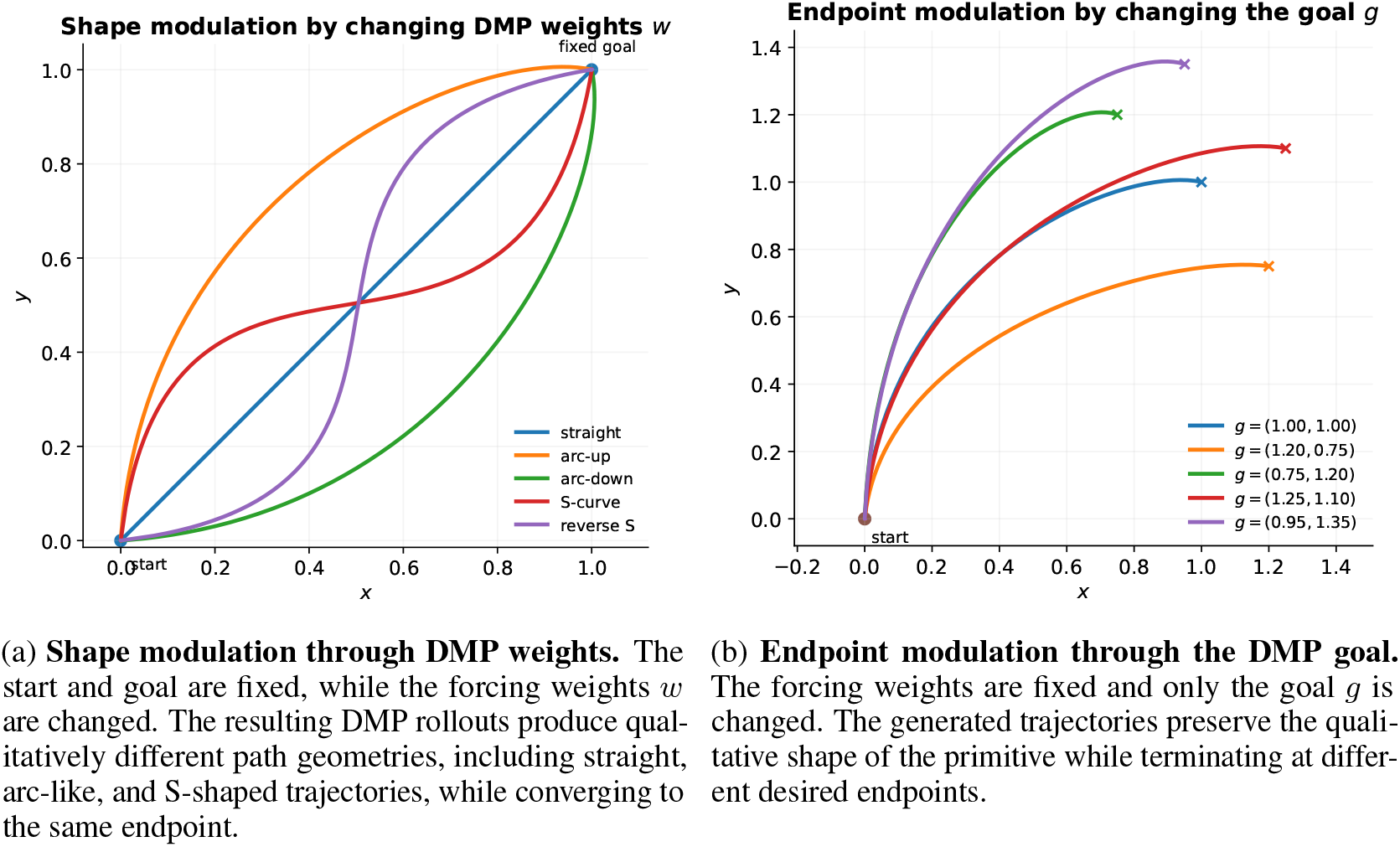
Controlled modulation properties of DMP trajectory generation. Left: varying the forcing weights *w* changes the trajectory shape while preserving the same start and goal. Right: varying the goal *g* changes the trajectory endpoint while preserving the qualitative trajectory structure.

#### Endpoint modulation by *g*

We next fixed the same forcing weights and changed only the goal *g*. Figure 5b shows that the rollout adapts to different terminal targets while preserving the same qualitative movement primitive. Across five goals, the mean terminal error was 4.71 × 10^−4^ and the maximum terminal error was 5.71 × 10^−4^. Thus, *g* provides direct endpoint control, whereas *w* provides shape control.

#### Autonomous recovery under perturbations

We then tested whether the DMP dynamics recover from mid-course state perturbations. A perturbation was applied at 45% of the movement duration, after which the DMP continued its rollout without replanning. Figure 6a shows representative recovery trajectories. The perturbed rollouts bend back toward the nominal movement and still converge to the goal.

**Figure 6:**
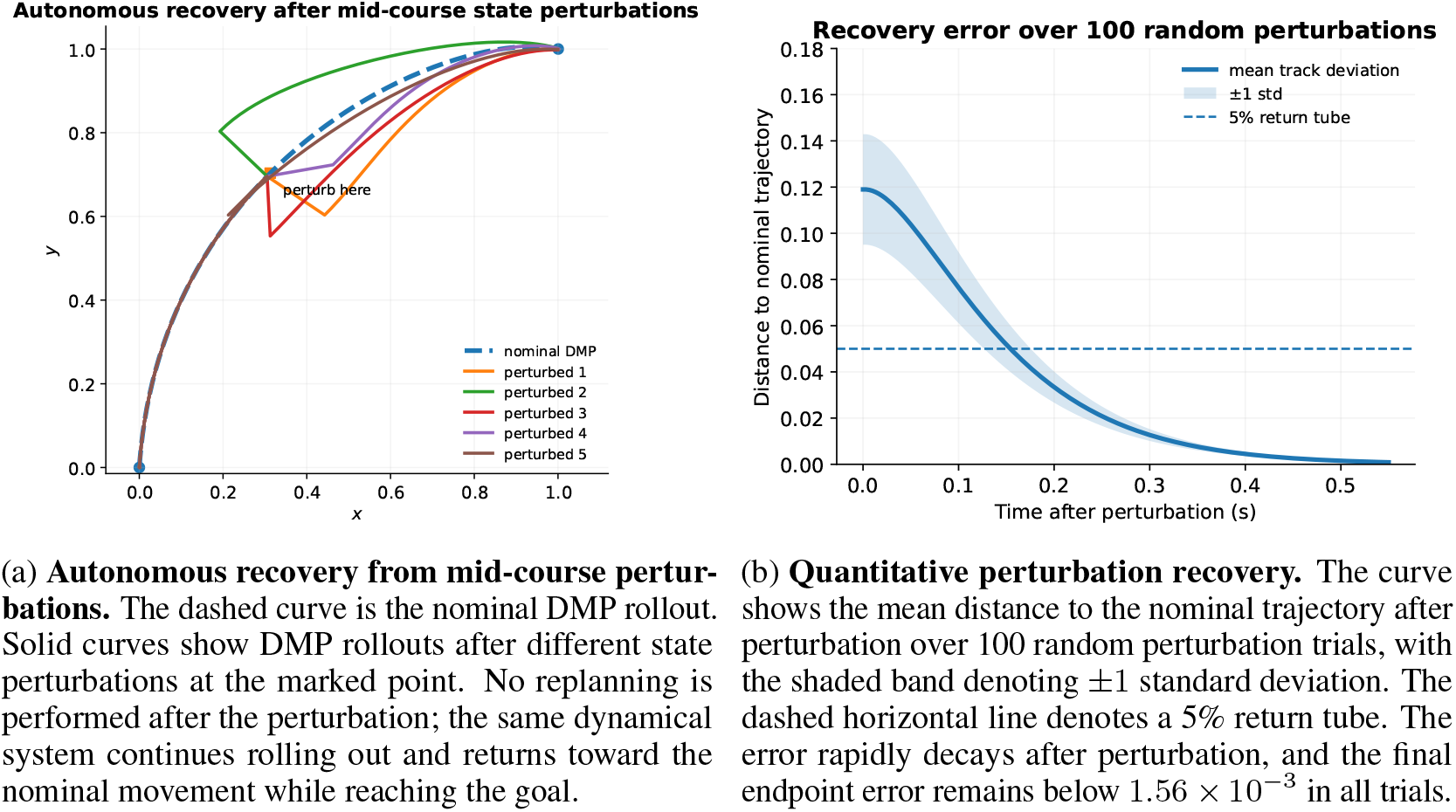
Autonomous perturbation recovery in DMP trajectory generation. Left: after mid-course state perturbations, the same DMP dynamics continue rolling out and return toward the nominal movement while still reaching the goal. Right: quantitative recovery analysis over 100 random perturbation trials shows that the deviation from the nominal trajectory decays rapidly after perturbation, with very small final endpoint error.

We further evaluated 100 random perturbations with directions sampled uniformly on [0, 2*π*) and magnitudes sampled uniformly from [0.08, 0.16](Figure 6b). The mean suffix deviation from the nominal trajectory was 0.0343 ± 0.0069. The endpoint error after perturbation was 9.23 × 10^−4^ on average and at most 1.56 × 10^−3^. Using a 5% return tube around the nominal trajectory, the mean return time was 0.153 ± 0.025 s, with median 0.157 s. These results show that the DMP generator is not merely an open-loop curve interpolator: the attractor dynamics provide autonomous correction after state disturbances.

#### Smooth point-to-point kinematics

Finally, we examined the time course of a DMP-generated point-to-point movement. We fitted a DMP to a smooth accelerate–cruise–decelerate reference profile and rolled it out using the same second-order generator. Figure 7 plots normalized position, speed, and acceleration. The DMP rollout preserves a smooth kinematic profile: position evolves monotonically, speed rises from rest, remains near its high-speed phase, and then decays smoothly to rest, while acceleration remains bounded and changes sign between the acceleration and deceleration phases. The rollout RMSE relative to the reference profile was 0.0031, and the endpoint error was 1.06 × 10^−3^. The speed remained above 90% of its peak for 0.593 s, illustrating the high-speed middle phase between acceleration and deceleration.

**Figure 7:**
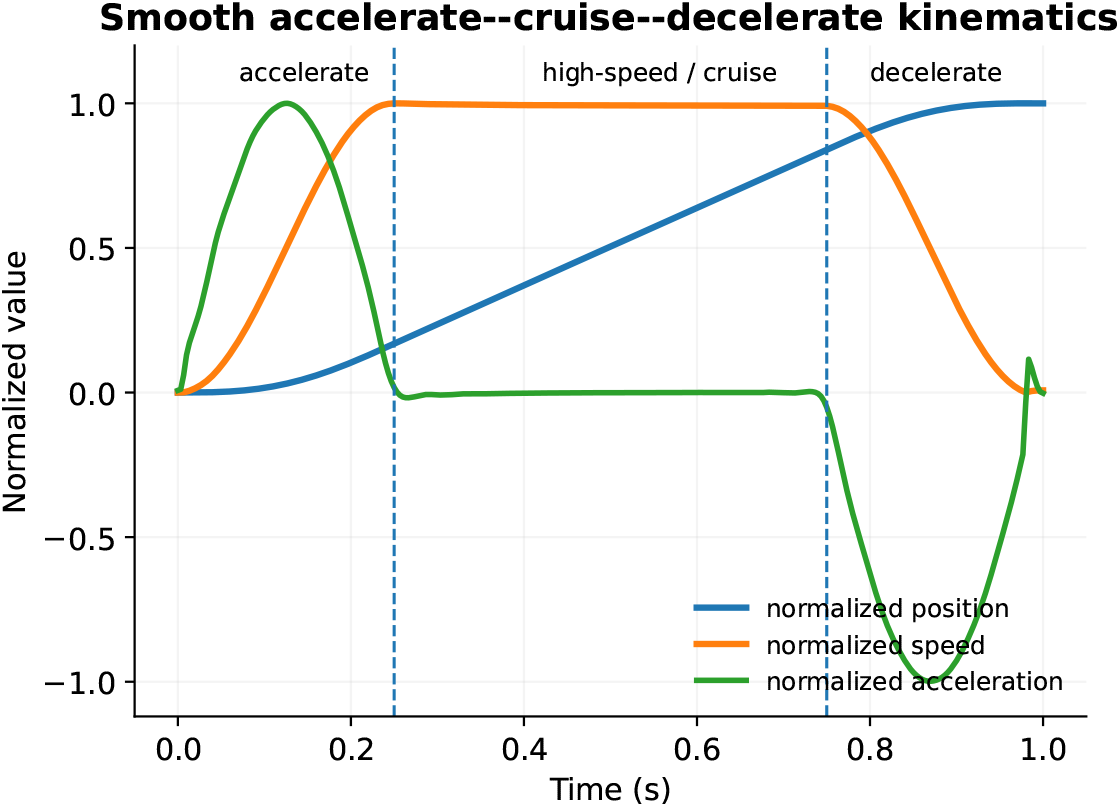
Smooth point-to-point kinematic profile. Normalized position, speed, and acceleration of a DMP-generated movement. The rollout exhibits a smooth accelerate–high-speed–decelerate structure: speed increases from rest, remains near its peak during the middle phase, and decreases smoothly near the end. This qualitative kinematic pattern is consistent with smooth biological point-to-point movement and avoids abrupt, temporally inconsistent motion.

#### Takeaway

These controlled experiments support the use of DMP as the output-side generator in Neural SS-DMP. Changing *w* modulates trajectory shape, changing *g* controls the endpoint, the second-order attractor recovers from state perturbations without explicit replanning, and the rollout produces smooth point-to-point kinematics. Therefore, DMP is not used merely as a smoother; it provides a structured movement generator that combines shape flexibility, goal-directed stability, and biologically plausible temporal organization.

### J.4 Sequential no-reset evaluation on M2

The main within-window protocol evaluates non-overlapping target segments independently. This is appropriate for controlled offline comparison, but it may overestimate performance for methods that rely on resetting each segment from target-aligned information. To probe whether the advantage of Neural SS-DMP persists under a more continuous rollout, we additionally evaluate a sequential no-reset setting on M2.

In this evaluation, consecutive non-overlapping held-out windows are processed in chronological order. Predictions are propagated across adjacent target segments rather than being reinitialized from Encoder output at each segment boundary. Thus, errors can accumulate over time, making this setting stricter than the independently reset within-window evaluation in Tab. 2. This experiment is still offline and does not constitute closed-loop or co-adaptive BCI validation; it is intended only as a stronger online-like stress test of continuous rollout behavior.

Table 8 shows that all methods degrade relative to the reset within-window evaluation. Nevertheless, Neural SS-DMP remains strongest in no-reset *R*^2^ and retains the lowest trajectory roughness. Although Kalman has a smaller absolute *R*^2^ drop, its accuracy remains substantially below Neural SS-DMP in both reset and no-reset settings. These results suggest that the structured-dynamics advantage is not limited to independently reset offline windows, while leaving true closed-loop and co-adaptive validation for future work.

**Table 8:** Sequential no-reset evaluation on M2. Variance-weighted *R*^2^ and mean squared jerk (MSJ; lower is better) are reported as mean ± 95% CI over 5 seeds. The reset *R*^2^ column repeats the corresponding M2 values from Tab. 2 for context. In the no-reset setting, predictions are propagated across consecutive held-out segments rather than being independently reinitialized at each target window.

| Method | Reset $R^2$ | No-reset $R^2$ | $\Delta R^2$ | No-reset MSJ |
| --- | --- | --- | --- | --- |
| Neural SS-DMP | **0.44 ± 0.08** | **0.40 ± 0.09** | −0.04 | **1.89 ± 0.14** |
| Transformer | 0.31 ± 0.10 | 0.22 ± 0.06 | −0.09 | 6.42 ± 0.58 |
| Kalman | 0.26 ± 0.07 | 0.23 ± 0.05 | −0.03 | 3.87 ± 0.22 |
| NoMAD | 0.41 ± 0.09 | 0.34 ± 0.07 | −0.07 | 3.21 ± 0.28 |

**Table 9:** Per-dimension metrics (Neural SS-DMP). Per-output-channel *R*^2^, correlation, and RMSE on the same evaluation splits as Tab. 2. Entries are mean ±95% CI over 5 seeds.

| Dataset | Dim | $R^2 \uparrow$ | Corr ↑ | RMSE ↓ |
| --- | --- | --- | --- | --- |
| M2 (spikes; $D=2$) | 1 | $0.43 \pm 0.09$ | $0.62 \pm 0.10$ | $0.72 \pm 0.16$ |
| | 2 | $0.42 \pm 0.08$ | $0.61 \pm 0.09$ | $0.74 \pm 0.15$ |
| H1 (spikes; $D=7$) | 1 | $0.22 \pm 0.06$ | $0.38 \pm 0.08$ | $1.21 \pm 0.14$ |
| | 2 | $0.18 \pm 0.07$ | $0.32 \pm 0.09$ | $1.35 \pm 0.18$ |
| | 3 | $0.20 \pm 0.06$ | $0.35 \pm 0.08$ | $1.29 \pm 0.16$ |
| | 4 | $0.25 \pm 0.05$ | $0.42 \pm 0.07$ | $1.15 \pm 0.15$ |
| | 5 | $0.35 \pm 0.05$ | $0.52 \pm 0.06$ | $0.98 \pm 0.11$ |
| | 6 | $0.31 \pm 0.05$ | $0.49 \pm 0.06$ | $1.06 \pm 0.13$ |
| | 7 | $0.31 \pm 0.06$ | $0.48 \pm 0.07$ | $1.07 \pm 0.14$ |
| BCI IV-4 (ECoG; $D=5$) | 1 | $0.42 \pm 0.09$ | $0.63 \pm 0.08$ | $0.74 \pm 0.10$ |
| | 2 | $0.38 \pm 0.08$ | $0.58 \pm 0.09$ | $0.79 \pm 0.09$ |
| | 3 | $0.32 \pm 0.09$ | $0.53 \pm 0.10$ | $0.86 \pm 0.11$ |
| | 4 | $0.26 \pm 0.10$ | $0.47 \pm 0.12$ | $0.94 \pm 0.13$ |
| | 5 | $0.22 \pm 0.11$ | $0.42 \pm 0.13$ | $1.01 \pm 0.15$ |
| MouseTrack (ECoG; $D=2$) | 1 | $0.35 \pm 0.10$ | $0.57 \pm 0.13$ | $0.76 \pm 0.09$ |
| | 2 | $0.49 \pm 0.07$ | $0.69 \pm 0.11$ | $0.68 \pm 0.08$ |

### J.5 Per-Dimension Metrics and Error Profiles

To complement the aggregate scores in Tab. 2, we report per-dimension *R*^2^, Pearson correlation (Corr), and RMSE for **Neural SS-DMP** on the same evaluation splits. Tab. 9 summarizes an error profile across degrees of freedom, indicating whether performance is broadly distributed across kinematic channels or concentrated in a subset. All values are mean ± 95% confidence interval over 5 seeds (computed with a *t*-interval across seeds).

### J.6 History length sweep (*H*; fixed *T* =0.5 s)

We sweep the causal history length *H* while keeping the decoded segment length fixed (*T* =0.5 s) to assess sensitivity to temporal context. Tab. 10 reports variance-weighted *R*^2^ across *H* on the same evaluation splits as Tab. 2. This sweep provides a compact ablation for the history window hyperparameter and checks that the default setting used in the main experiments lies in a stable regime. Across datasets, *H*=0.7 s is best or near-best, supporting the default setting used in Tab. 2. Increasing history beyond 0.7 s does not consistently help: *H*=1.0 s yields small degradations across all datasets, indicating diminishing returns under the fixed-*T* protocol.

### J.7 Learned Subject-Dynamics Priors: Three Focused Analyses

We focus on three questions: (1) how learned priors differ from the DMP base, (2) how learned priors differ across datasets, and (3) how learned priors vary across output dimensions within a dataset.

#### Prior parameterization

For each output dimension, we fit a continuous-time second-order linear prior

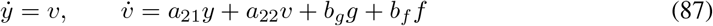

equivalently 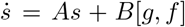 with 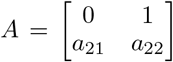 and 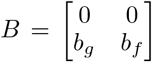. We fit (*A*_*d*_, *B*_*d*_) per output dimension *d*.Reported means and standard deviations are computed across output dimensions within each dataset.

1. **Learned vs. base**. Figure 8 shows the dataset-level mean matrices (*A*_learned_, *B*_learned_), and Table 11 reports the corresponding coefficients. These summaries make explicit how the fitted linear dynamics compare to the shared DMP base. Across datasets, the learned dynamics preserve the canonical state-space structure (*a*_12_=1, zeros elsewhere) while adjusting the second-order coefficients: *a*_21_ and *a*_22_ remain negative (stable attractor form) but shift in magnitude in a dataset-dependent way, indicating different effective stiffness/damping relative to the base. The learned input gains also change systematically: *b*_*g*_ tracks the scale of |*a*_21_| (consistent with how the goal term enters the acceleration dynamics), while *b*_*f*_ deviates from the base value of 1 and varies across datasets, reflecting dataset-specific rescaling of the forcing pathway. Notably, these differences are observed under a matched horizon duration (*T* Δ*t*=0.5 s), so the coefficient shifts reflect learned dynamics rather than changes in rollout duration.
2. **Cross-dataset differences**. Figure 9 visualizes the dataset-level coefficient means (bars), with gray dashed lines indicating the base DMP coefficients. Relative to the base, *a*_21_ becomes more negative for H1 and both ECoG datasets (stronger restoring term), while M2 remains closer to the base; *a*_22_ also shifts in magnitude, with the largest damping observed on BCI IV-4. The input gains change in tandem: *b*_*g*_ increases when |*a*_21_| increases (consistent with the goal term scaling), and *b*_*f*_ is systematically above the base value of 1 for all but M2, indicating a dataset-dependent rescaling of the forcing pathway under the same matched horizon duration (*T* Δ*t*=0.5 s).
3. **Within-dataset variability across output dimensions**. Figure 10 summarizes per-dimension variability within each dataset, with red dashed lines marking the base coefficients. The 2D tasks (M2 and MouseTrack) show tight distributions across dimensions, indicating relatively homogeneous fitted dynamics. In contrast, the higher-DoF datasets (H1 and BCI IV-4) exhibit noticeably larger dispersion—most prominently in *a*_21_ (restoring strength) and *b*_*g*_/*b*_*f*_ (input gains)—suggesting that different output channels favor different effective stiffness/gain settings even under the shared stable second-order form.

**Table 10:**
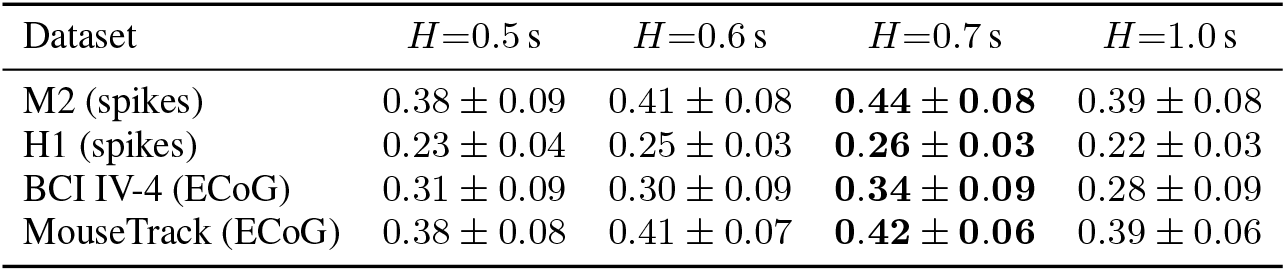
History length sweep. Variance-weighted *R*^2^ (higher is better). Values at *H*=0.7 s match Tab. 2.

| Dataset | $H=0.5$ s | $H=0.6$ s | $H=0.7$ s | $H=1.0$ s |
| --- | --- | --- | --- | --- |
| M2 (spikes) | $0.38 \pm 0.09$ | $0.41 \pm 0.08$ | **$0.44 \pm 0.08$** | $0.39 \pm 0.08$ |
| H1 (spikes) | $0.23 \pm 0.04$ | $0.25 \pm 0.03$ | **$0.26 \pm 0.03$** | $0.22 \pm 0.03$ |
| BCI IV-4 (ECoG) | $0.31 \pm 0.09$ | $0.30 \pm 0.09$ | **$0.34 \pm 0.09$** | $0.28 \pm 0.09$ |
| MouseTrack (ECoG) | $0.38 \pm 0.08$ | $0.41 \pm 0.07$ | **$0.42 \pm 0.06$** | $0.39 \pm 0.06$ |

**Table 11:** Numeric summary of subject-dynamics prior coefficients in Eq. (87). Δ*t* is the sampling interval, *T* is the rollout horizon in *steps*, and *D* is the output dimensionality. Values are mean ± std across output dimensions within each dataset. All datasets use a matched horizon duration (*T* · Δ*t* = 0.5 s), consistent with the main-text decoding horizon.

| Dataset | $\Delta t$ (s) | $T$ | $D$ | $a_{21}$ | $a_{22}$ | $b_g$ | $b_f$ |
| --- | --- | --- | --- | --- | --- | --- | --- |
| Base (DMP) | — | — | — | -156.25 | -25.00 | 156.25 | 1.000 |
| M2 (spikes) | 0.020 | 25 | 2 | -136.11±0.49 | -19.42±0.15 | 133.90±0.61 | 0.860±0.006 |
| H1 (spikes) | 0.020 | 25 | 7 | -210.86±7.00 | -32.70±0.98 | 210.88±7.00 | 1.396±0.050 |
| BCI IV-4 (ECoG) | 0.001 | 500 | 5 | -283.89±7.99 | -42.43±1.59 | 270.17±3.31 | 1.584±0.068 |
| MouseTrack (ECoG) | 0.001 | 500 | 2 | -278.09±7.64 | -35.90±2.02 | 276.15±6.80 | 1.729±0.075 |

**Figure 8:**
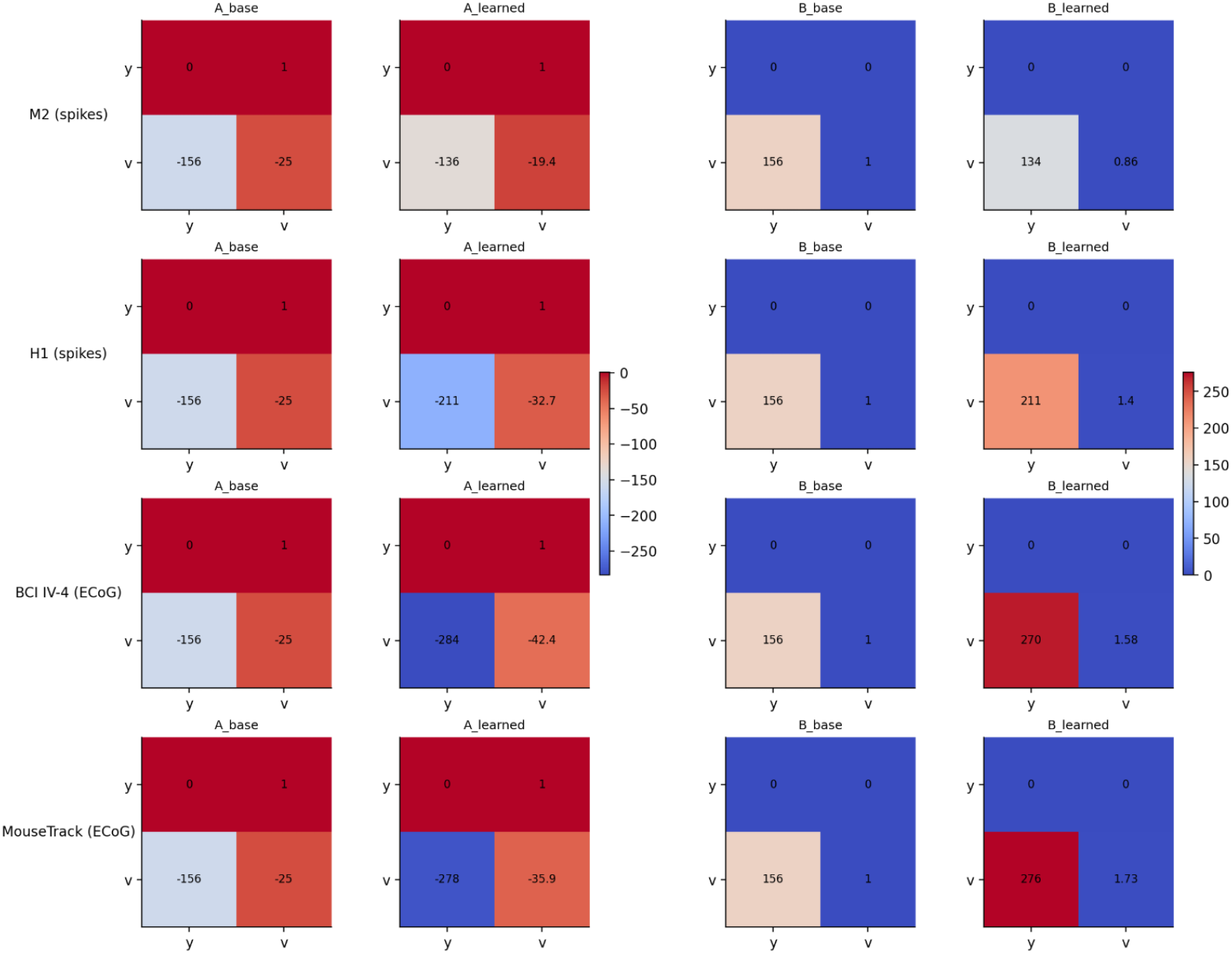
Dataset-level mean matrices. For each dataset, *A*_learned_ and *B*_learned_ are obtained by averaging the fitted per-dimension matrices (*A*_*d*_, *B*_*d*_) across output dimensions; *A*_base_ and *B*_base_ are the shared DMP base matrices.

**Figure 9:**
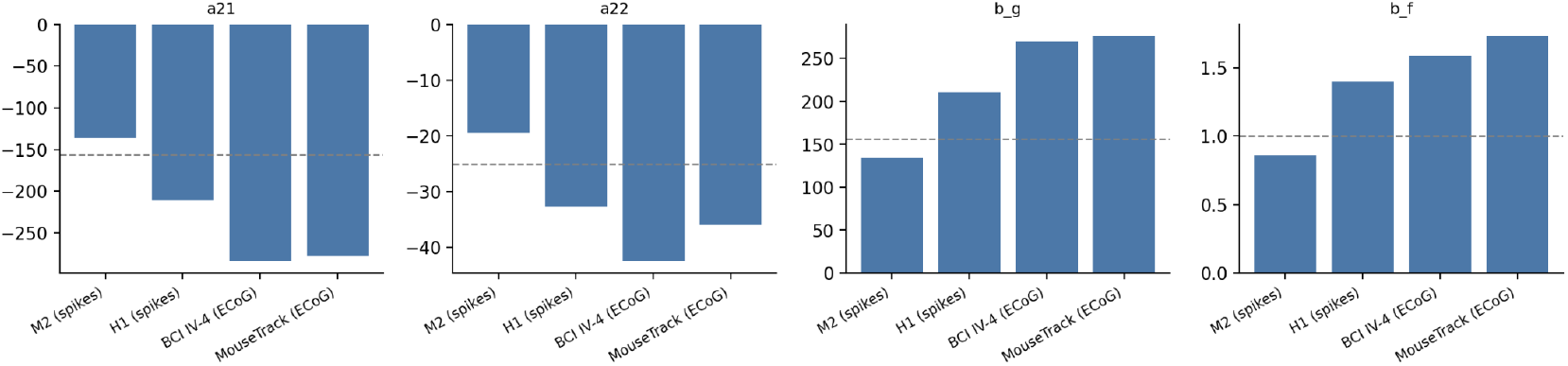
Cross-dataset comparison of learned coefficient means (bars). Gray dashed lines indicate the base DMP coefficients.

**Figure 10:**
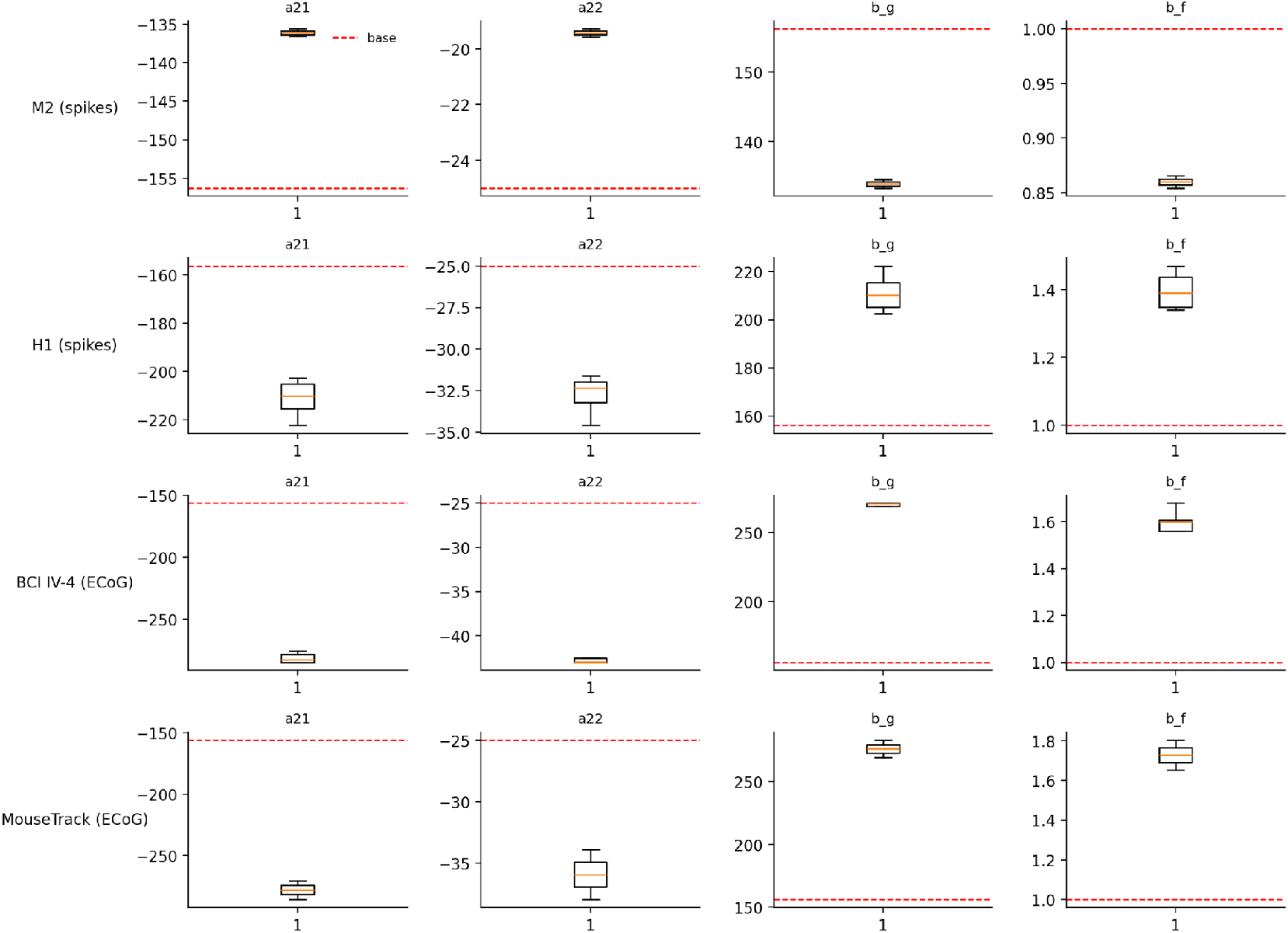
Within-dataset variability across output dimensions. Each box summarizes the distribution of per-dimension fitted coefficients; red dashed lines indicate base coefficients.

### J.8 ECoG feature ablation: raw vs. PSD vs. raw+PSD fusion

We ablate the ECoG input representation by training the same architecture under identical evaluation splits and training budgets with (i) raw time-domain features only, (ii) PSD (band-power) features only, and (iii) raw+PSD multimodal fusion (main setting). Tab. 12 reports within-window performance (*H*=0.7 s, *T* =0.5 s) on the ECoG benchmarks. Fusion performs best across metrics, suggesting that time-domain and spectral cues provide complementary information for decoding.

### J.9 Prior-strength extensions: joint sweep over *λ*_*A*_ **and** *λ*_*B*_

We extend the main-text prior-strength ablation (Sec. 4.4.4) by sweeping *separately* the blend applied to the state matrix (*λ*_*A*_) and the input matrix (*λ*_*B*_). Concretely, when subject priors are enabled, we use

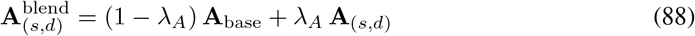

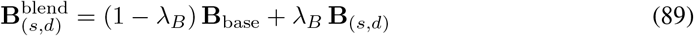

and evaluate the resulting decoder under the *same* windowing protocol, splits, and metrics as the main results (Tab. 2). We perform a *joint* grid search over (*λ*_*A*_, *λ*_*B*_) (rather than two independent 1D sweeps) to capture potential synergy between stabilizing the homogeneous dynamics (**A**) and calibrating the goal/forcing channels (**B**).

#### Evaluation protocol

The default prior strengths (*λ*_*A*_, *λ*_*B*_) = (0.2, 0.5) used in all main-text experiments were selected once using the held-out validation split before any test-set reporting. No test-set kinematics were used for fitting the prior, selecting (*λ*_*A*_, *λ*_*B*_), early stopping, calibration, or model selection. To assess robustness to the choice of prior strength, we additionally report test-set variance-weighted *R*^2^ on the same evaluation splits as Tab. 2 (HI → HO for spikes; subject-wise temporal test split for ECoG), aggregated across subjects/sessions per the main protocol. Entries are mean ± 95% CI over five random seeds. This sweep is reported as a sensitivity analysis only, not as the procedure used to choose the default hyperparameters.

#### Results and choice of default

Table 13 shows that moderate prior blending yields a stable high-performing regime across datasets. Performance is relatively flat around *λ*_*A*_ ∈ { 0.1, 0.2} and *λ*_*B*_ ∈ { 0.25, 0.5}, while overly large blends can over-constrain the dynamics or over-amplify the goal/forcing channels. The validation-selected default (*λ*_*A*_, *λ*_*B*_) = (0.2, 0.5) lies in this stable regime. We therefore interpret Table 13 as evidence of robustness to the prior-strength choice, not as test-set hyperparameter selection.

**Table 12:** ECoG feature ablation (Neural SS-DMP only). Within-window decoding (*H*=0.7 s, *T* =0.5 s). We compare raw-only, PSD-only, and raw+PSD fusion under identical splits and training budgets. Entries are mean ± 95% CI across *5* random seeds.

| Dataset | Metric | Raw only | PSD only | Raw+PSD |
| --- | --- | --- | --- | --- |
| BCI IV-4 | $R^2 \uparrow$ | $0.31 \pm 0.10$ | $0.28 \pm 0.11$ | **$0.34 \pm 0.09$** |
| | Corr ↑ | $0.50 \pm 0.12$ | $0.46 \pm 0.13$ | **$0.54 \pm 0.11$** |
| | RMSE ↓ | $0.86 \pm 0.08$ | $0.90 \pm 0.10$ | **$0.82 \pm 0.07$** |
| MouseTrack (avg.) | $R^2 \uparrow$ | $0.40 \pm 0.07$ | $0.37 \pm 0.08$ | **$0.42 \pm 0.06$** |
| | Corr ↑ | $0.61 \pm 0.11$ | $0.57 \pm 0.13$ | **$0.63 \pm 0.12$** |
| | RMSE ↓ | $0.75 \pm 0.09$ | $0.81 \pm 0.10$ | **$0.72 \pm 0.08$** |

**Table 13:** Joint prior-strength sweep over (*λ*_*A*_, *λ*_*B*_). Test-set variance-weighted *R*^2^ (mean ± 95% CI over 5 seeds). We jointly scan *λ*_*A*_ ∈ { 0, 0.1, 0.2, 0.3, 0.5} and *λ*_*B*_ ∈ {0, 0.25, 0.5, 0.75, 1.0} . Best observed value in each dataset block is bolded for orientation only.

| M2 (spikes; $D=2$) | | | | | | H1 (spikes; $D=7$) | | | | | |
| --- | --- | --- | --- | --- | --- | --- | --- | --- | --- | --- | --- |
| $\lambda_A \backslash \lambda_B$ | 0 | 0.25 | 0.5 | 0.75 | 1.0 | $\lambda_A \backslash \lambda_B$ | 0 | 0.25 | 0.5 | 0.75 | 1.0 |
| 0.0 | 0.37 ± 0.09 | 0.39 ± 0.08 | 0.42 ± 0.08 | 0.38 ± 0.09 | 0.34 ± 0.10 | 0.0 | 0.24 ± 0.04 | 0.24 ± 0.04 | 0.25 ± 0.03 | 0.23 ± 0.04 | 0.21 ± 0.05 |
| 0.1 | 0.41 ± 0.08 | 0.42 ± 0.07 | 0.43 ± 0.08 | 0.40 ± 0.09 | 0.36 ± 0.11 | 0.1 | 0.24 ± 0.04 | 0.25 ± 0.03 | 0.25 ± 0.03 | 0.24 ± 0.04 | 0.22 ± 0.05 |
| 0.2 | 0.43 ± 0.09 | 0.44 ± 0.08 | **0.44 ± 0.08** | 0.41 ± 0.09 | 0.37 ± 0.10 | 0.2 | 0.25 ± 0.04 | 0.26 ± 0.03 | **0.26 ± 0.03** | 0.24 ± 0.04 | 0.22 ± 0.04 |
| 0.3 | 0.41 ± 0.10 | 0.43 ± 0.09 | 0.43 ± 0.09 | 0.39 ± 0.10 | 0.35 ± 0.11 | 0.3 | 0.24 ± 0.04 | 0.25 ± 0.04 | 0.25 ± 0.04 | 0.23 ± 0.04 | 0.20 ± 0.05 |
| 0.5 | 0.36 ± 0.11 | 0.38 ± 0.10 | 0.39 ± 0.10 | 0.35 ± 0.12 | 0.31 ± 0.13 | 0.5 | 0.21 ± 0.05 | 0.22 ± 0.04 | 0.23 ± 0.04 | 0.21 ± 0.05 | 0.18 ± 0.06 |

| BCI IV-4 (ECoG; $D=5$) | | | | | | MouseTrack (ECoG; $D=2$) | | | | | |
| --- | --- | --- | --- | --- | --- | --- | --- | --- | --- | --- | --- |
| $\lambda_A \backslash \lambda_B$ | 0 | 0.25 | 0.5 | 0.75 | 1.0 | $\lambda_A \backslash \lambda_B$ | 0 | 0.25 | 0.5 | 0.75 | 1.0 |
| 0.0 | 0.33 ± 0.09 | 0.33 ± 0.08 | 0.34 ± 0.09 | 0.32 ± 0.09 | 0.30 ± 0.10 | 0.0 | 0.40 ± 0.07 | 0.40 ± 0.07 | 0.41 ± 0.06 | 0.40 ± 0.07 | 0.38 ± 0.08 |
| 0.1 | 0.33 ± 0.08 | 0.34 ± 0.08 | 0.34 ± 0.08 | 0.33 ± 0.09 | 0.31 ± 0.10 | 0.1 | 0.41 ± 0.06 | 0.41 ± 0.06 | 0.42 ± 0.06 | 0.41 ± 0.07 | 0.39 ± 0.08 |
| 0.2 | 0.33 ± 0.09 | 0.34 ± 0.08 | **0.34 ± 0.09** | 0.32 ± 0.09 | 0.30 ± 0.11 | 0.2 | 0.41 ± 0.07 | 0.42 ± 0.06 | **0.42 ± 0.06** | 0.40 ± 0.07 | 0.38 ± 0.08 |
| 0.3 | 0.32 ± 0.09 | 0.33 ± 0.09 | 0.33 ± 0.09 | 0.31 ± 0.10 | 0.29 ± 0.11 | 0.3 | 0.40 ± 0.07 | 0.41 ± 0.07 | 0.41 ± 0.07 | 0.39 ± 0.08 | 0.37 ± 0.09 |
| 0.5 | 0.30 ± 0.10 | 0.31 ± 0.10 | 0.31 ± 0.10 | 0.29 ± 0.11 | 0.27 ± 0.12 | 0.5 | 0.37 ± 0.08 | 0.38 ± 0.08 | 0.39 ± 0.07 | 0.37 ± 0.09 | 0.34 ± 0.10 |

### J.10 Velocity-distribution shift stress test on M2

A potential limitation of behavior-prior decoders is that a prior fit from training kinematics may become less useful when test movements have different speed statistics. We therefore evaluate a velocity-distribution shift stress test on M2. For each segment, we compute the scalar speed profile *v*_*t*_ = ∥**y**_*t*+1_ − **y**_*t*_ ∥_2_*/*Δ*t* and summarize its distribution using mean speed, peak speed, speed standard deviation, and empirical speed quantiles. Using training segments only, we estimate the training speed-distribution statistics. Each held-out test segment is assigned a velocity-shift score by its standardized distance from the training speed-feature distribution, and held-out windows are partitioned into low-, medium-, and high-shift bins by tertiles of this score.

This analysis uses test kinematics only for post-hoc evaluation stratification, not for model fitting, hyperparameter selection, or recalibration. It asks whether the behavior-derived prior remains useful when held-out movements differ from the training speed distribution. Table 14 reports the resulting *R*^2^ and MSJ across shift bins.

**Table 14:**
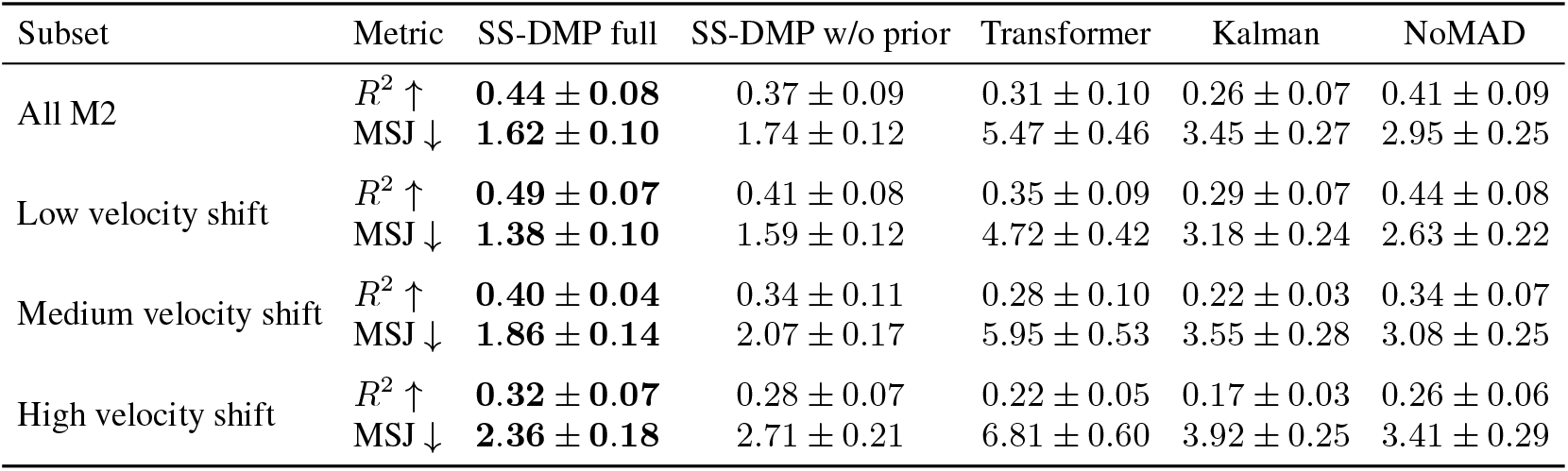
Velocity-distribution shift stress test on M2. Held-out M2 windows are stratified into low-, medium-, and high-shift bins according to the distance of their speed-distribution features from the training speed distribution. We report variance-weighted *R*^2^ and mean squared jerk (MSJ). The “All M2” row reproduces the corresponding M2 results from the main tables for reference.

| Subset | Metric | SS-DMP full | SS-DMP w/o prior | Transformer | Kalman | NoMAD |
| --- | --- | --- | --- | --- | --- | --- |
| All M2 | $R^2 \uparrow$ | **0.44 ± 0.08** | 0.37 ± 0.09 | 0.31 ± 0.10 | 0.26 ± 0.07 | 0.41 ± 0.09 |
| | MSJ ↓ | **1.62 ± 0.10** | 1.74 ± 0.12 | 5.47 ± 0.46 | 3.45 ± 0.27 | 2.95 ± 0.25 |
| Low velocity shift | $R^2 \uparrow$ | **0.49 ± 0.07** | 0.41 ± 0.08 | 0.35 ± 0.09 | 0.29 ± 0.07 | 0.44 ± 0.08 |
| | MSJ ↓ | **1.38 ± 0.10** | 1.59 ± 0.12 | 4.72 ± 0.42 | 3.18 ± 0.24 | 2.63 ± 0.22 |
| Medium velocity shift | $R^2 \uparrow$ | **0.40 ± 0.04** | 0.34 ± 0.11 | 0.28 ± 0.10 | 0.22 ± 0.03 | 0.34 ± 0.07 |
| | MSJ ↓ | **1.86 ± 0.14** | 2.07 ± 0.17 | 5.95 ± 0.53 | 3.55 ± 0.28 | 3.08 ± 0.25 |
| High velocity shift | $R^2 \uparrow$ | **0.32 ± 0.07** | 0.28 ± 0.07 | 0.22 ± 0.05 | 0.17 ± 0.03 | 0.26 ± 0.06 |
| | MSJ ↓ | **2.36 ± 0.18** | 2.71 ± 0.21 | 6.81 ± 0.60 | 3.92 ± 0.25 | 3.41 ± 0.29 |

**Table 15:** Quantifying forcing-geometry structure in 3D PCA space. Fits are computed on the 3D PCA scores underlying Fig. 11. BCI IV-4 is summarized by a best-fit plane explained fraction; H1 is summarized by a union-of-rays model with selected *K*_ray_.

| Dataset | $N$ | Model | Explained | Notes |
| --- | --- | --- | --- | --- |
| BCI IV-4 (ECoG) | 800 | plane (+ thin ridge) | 0.9989 | near rank-2 with a line-like excursion |
| H1 (spikes) | 2500 | union of rays | 0.7885 | $K_{\text{ray}} = 6$ rays; branches through a hub |

#### Takeaway

As velocity-distribution shift increases, all methods degrade, indicating that high-shift test movements are genuinely harder. The advantage of the full SS-DMP model over the no-prior variant narrows in the high-shift bin, which is consistent with the learned behavior prior being most useful when test speeds resemble the training speed distribution. However, even under high velocity shift, the full model remains competitive in *R*^2^ and preserves the best MSJ among the compared methods. Thus, the learned behavior prior is not harmful in this stress setting, but its benefit should be understood as a structured inductive bias for matched or moderately shifted continuous movement regimes rather than a universal guarantee for arbitrary speed distributions.

## K Interpretability: Forcing Geometry

This appendix analyzes the geometry of the predicted forcing input at the SS-DMP generator. The goal is to determine whether the encoder sends arbitrary high-dimensional sequence outputs into the generator, or whether the predicted forcing trajectories occupy a compact and structured control space. We focus on **BCI IV-4** (ECoG) and **H1** (spikes), where the forcing embeddings show stable low-dimensional structure.

### K.1 Construction of Forcing Embeddings

#### Model-predicted forcing

For each evaluation window, we compute the *predicted* forcing trajectory 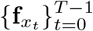 induced by the model outputs (**ŵ**, **ĝ, Ŷ**_0_) through the SS-DMP forcing function in Eq. (3).

Here 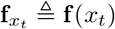 denotes the discretized forcing at DMP phase *x*_*t*_.

#### Phase/amplitude normalization

To factor out trivial phase and amplitude scaling, we analyze the normalized forcing

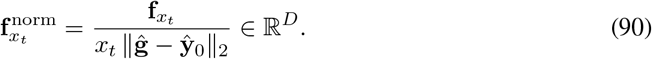

This normalization removes the explicit phase factor *x*_*t*_ and the predicted displacement scale ∥**ĝ** − **Ŷ**_0_∥_2_, so the PCA primarily reflects forcing-shape variability rather than trivial movement amplitude or phase decay.

#### Prefix restriction and vectorization

Because *x*_*t*_ → 0 near termination, the normalization in Eq. (90) can amplify late-bin numerical noise. We therefore restrict analysis to a prefix where *x*_*t*_ is bounded away from zero and define

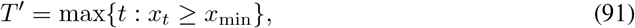

equivalently keeping *t* = 0, …, *T* ^*′*^ − 1, with a fixed threshold *x*_min_ shared across windows within each dataset. We then stack the normalized forcing over this prefix into one vector

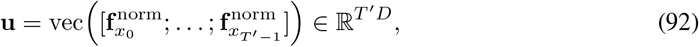

and fit PCA across evaluation windows within each dataset.

Although **u** is represented in ℝ^*T*′*D*^, the forcing trajectory is functionally constrained by the RBF basis in Eq. (3): each **ŵ** _*d*_ lies in the span of *K* basis functions, so temporal variation lives in a low-dimensional function subspace with on the order of *DK* degrees of freedom, not unconstrained *T* ^*′*^*D* degrees of freedom.

#### Window sampling, axes, and color

All points are sampled from the same evaluation split used by the corresponding main results. For consistent rendering, we cap the number of plotted windows per dataset and subsample with a fixed seed: BCI IV-4 uses *N* = 800 windows and H1 uses *N* = 2500 windows. To prevent a small number of outliers from dominating scatter-axis scaling, axis limits are set by the 2nd–98th percentiles within each dataset. Point color indicates predicted displacement amplitude ∥**ĝ** − **Ŷ**_0_∥_2_ for reference.

### K.2 Compact Forcing Geometry at the Generator Input

#### Forcing variability concentrates into few modes

Across evaluation windows, normalized forcing-shape variability concentrates into only a few principal components (Fig. 11, top). For BCI IV-4 (*D* = 5), the first five PCs explain 0.71 of the variance; for H1 (*D* = 7), the first seven PCs explain 0.92 of the variance. This suggests that the encoder maps diverse neural histories into a compact set of forcing-shape modes compatible with stable rollout. This is not a claim that the kinematics are intrinsically low-dimensional; rather, it characterizes the learned generator input after normalization and SS-DMP parameterization.

**Figure 11:**
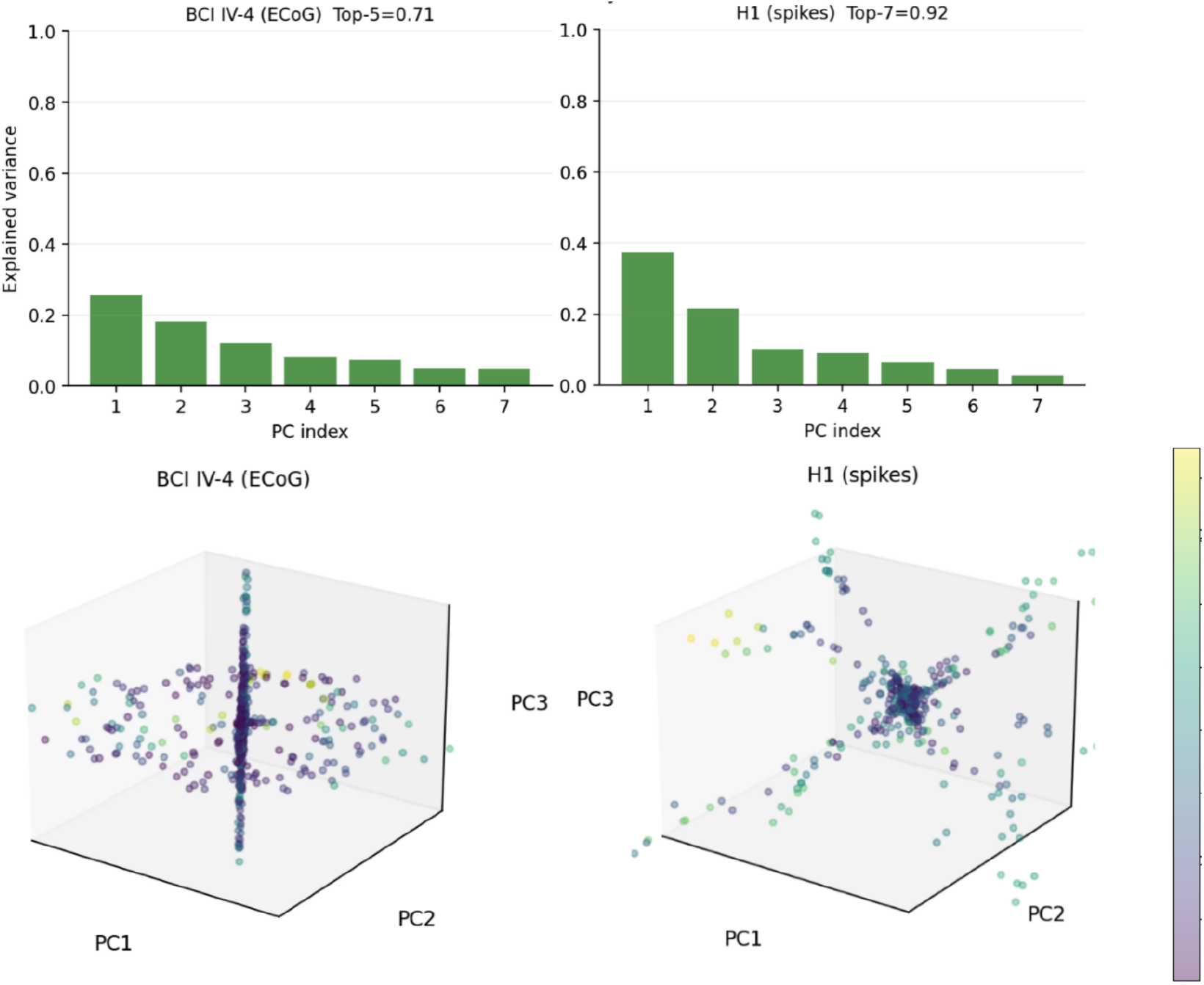
Compact forcing geometry at the generator input (BCI IV-4 and H1). **Top:** PCA explained-variance ratios of 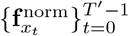 (Eq. (90)) across evaluation windows, computed separately per dataset. Titles report Top-*D*, the cumulative variance explained by the first *D* PCs, where *D* equals the kinematic dimension. **Bottom:** 3D PCA scatter of the normalized forcing vectors across windows. Points are colored by predicted displacement amplitude ∥**ĝ** − **Ŷ**_0_∥_2_.

#### Geometry suggests discrete regimes with continuous modulation

The leading 3D PCA embeddings are strongly anisotropic and dataset-specific (Fig. 11, bottom). BCI IV-4 concentrates near a low-dimensional continuum with a thin orthogonal component, whereas H1 exhibits a hub-and-branches pattern consistent with a union of low-dimensional subspaces. Under the SS-DMP view, this provides a mechanistic interpretation of robustness under an explicit dynamical prior: predicted forcing is organized into a small set of stable directions that the dynamics can reliably realize, rather than being absorbed into unconstrained high-dimensional sequence outputs.

### K.3 Quantifying Planes and Rays in 3D PCA Space

Visual inspection of Fig. 11 suggests distinct low-dimensional geometries: BCI IV-4 concentrates near a plane with an additional thin, line-like excursion, whereas H1 exhibits a multi-branch union-of-rays structure. We quantify these patterns directly in the 3D PCA score space. Let **z**_*n*_ ∈ ℝ^3^ denote the 3D PCA score of evaluation window *n*. Quantitative summaries are reported in Tab. 15.

#### BCI IV-4: best-fit plane with a line-like ridge

Let 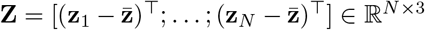 be the centered score matrix. We fit a plane by taking the top two right singular vectors of **Z** and let **P**_2_ denote the orthogonal projector onto this 2D subspace. We report the explained fraction

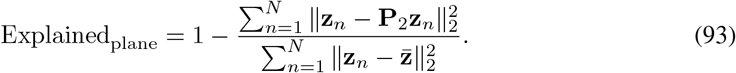

A value near one indicates an almost rank-2 embedding, consistent with the near-planar structure in Fig. 11 and the pairwise 2D projections in Fig. 12.

**Figure 12:**
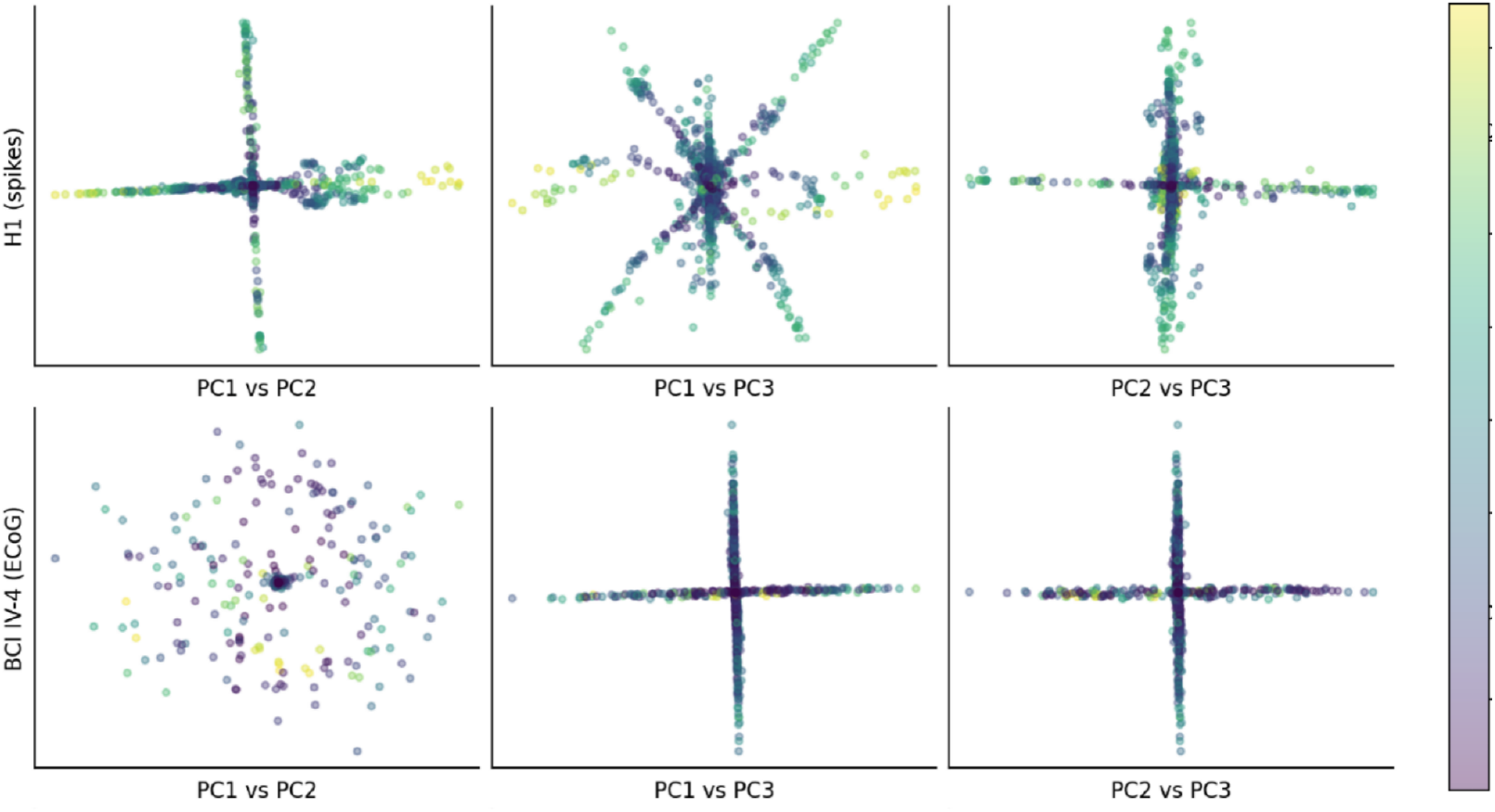
Pairwise 2D projections of forcing-input PCA scores. Rows correspond to datasets (H1, BCI IV-4). Columns show PC1–PC2, PC1–PC3, and PC2–PC3 projections of the same 3D PCA embedding shown in Fig. 11. Points are colored by predicted displacement amplitude ∥**ĝ** − **Ŷ**_0_∥_2_.

#### H1: union of rays through a shared hub

We model the H1 embedding as a union of *K*_ray_ rays through a shared hub 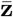. Let 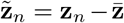 and 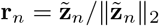 denote unit directions. We cluster **r**_*n*_ on the sphere using direction-only clustering with cosine distance and a fixed random seed, yielding ray directions 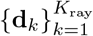. Each point is reconstructed by orthogonal projection onto its assigned ray:

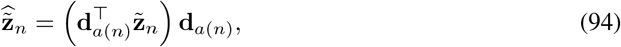

where *a*(*n*) is the assigned cluster. We report

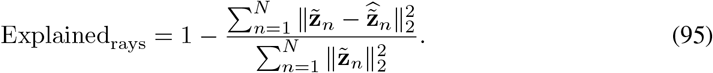

We select *K*_ray_ by an elbow criterion over *K*_ray_ ∈ {2, …, 6}, choosing the smallest value beyond which gains saturate.

### K.4 Print-Friendly 2D Projections

Figure 12 shows pairwise projections of the same 3D PCA embedding as Fig. 11. These views facilitate static PDF inspection and help verify that the structures are not artifacts of a single 3D viewing angle. BCI IV-4 remains dominated by a planar slab with a thin ridge through the hub, while H1 preserves a multi-branch star-like geometry across all projections.

### K.5 Interpretation

The forcing trajectory is the generator input that shapes motion beyond the stable second-order prior, so its embedding has a direct mechanistic interpretation: it describes the effective control space used by the model to generate movement within the SS-DMP family. In BCI IV-4, the near-planar embedding suggests that most forcing-shape variability is governed by a small number of smoothly varying control factors, with an additional thin mode capturing rarer deviations. In H1, a 7-DoF robotic-arm setting, the multi-ray structure is consistent with a small number of distinct control regimes that are continuously modulated within each regime. Together, these results suggest that the structured dynamical generator does not merely smooth predictions, but induces an interpretable and compact control space whose organization can be read out from neural-driven forcing.

